# Deep profiling of mouse splenic architecture with CODEX multiplexed imaging

**DOI:** 10.1101/203166

**Authors:** Yury Goltsev, Nikolay Samusik, Julia Kennedy-Darling, Salil Bhate, Matthew Hale, Gustavo Vazquez, Sarah Black, Garry P. Nolan

## Abstract

A highly multiplexed cytometric imaging approach, termed CO-Detection by indEXing (CODEX), is used here to create multiplexed datasets of normal and lupus (MRL/*lpr*) murine spleens. CODEX iteratively visualizes antibody binding events using DNA barcodes, fluorescent dNTP analogs, and an insitu polymerization-based indexing procedure. An algorithmic pipeline for single-cell antigen quantification in tightly packed tissues was developed and used to overlay well-known morphological features with *de novo* characterization of lymphoid tissue architecture at a single-cell and cellular neighborhood levels. We observed an unexpected, profound impact of the cellular neighborhood on the expression of protein receptors on immune cells. By comparing normal murine spleen to spleens from animals with systemic autoimmune disease (MRL/*lpr*), extensive and previously uncharacterized splenic cell interaction dynamics in the healthy versus diseased state was observed. The fidelity of multiplexed spatial cytometry demonstrated here allows for quantitative systemic characterization of tissue architecture in normal and clinically aberrant samples.

## Introduction

Dramatic immune tissue re-organization has been seen in lupus erythematosus, where a variety of organs (from skin, to kidney, and other body organs) can be targeted in relapsing-remitting flares. One example of such reorganization is pronounced lymphadenopathy and splenomegaly observed in lupus models (Lieberum and Hartmann, 1988, Jacobson et al., 1995). Using mice with MRL/*lpr* genotype (Kanauchi et al., 1991) we sought to systematically characterizes microenvironment and cell interactions associated with changes in immune organ architecture and the progression of autoimmune disease. To this end we devised a multiplexed microscopy technique that allows a precise mapping of cell types in tissues. Significant overlap in excitation and emission spectra makes it hard to image more than 4-5 fluorophores with conventional fluorescent microscopy. Yet considerably more surface markers are needed for precise identification of cellular subsets and their activation state (Chattopadhyay and Roederer, 2012). Approaches have been developed to overcome such limitations (Schubert et al., 2006, Gerdes et al., 2013), but these protocols have required multiple stain/strip/wash cycles of the antibodies that can be time consuming or lead to sample degradation over the iterations.

The technique described here (CODEX, for <u>CO</u>-<u>D</u>etection by ind<u>EX</u>ing) extends deep phenotyping capabilities of flow and mass cytometry (Spitzer et al., 2015, Bendall et al., 2011) to most standard three-color fluorescence microscope platforms for imaging of solid tissues. Accurate highly multiplexed single-cell quantification of membrane protein expression in densely packed lymphoid tissue images, (which was once deemed impossible (Gerner et al., 2012)) was achieved using polymerase driven incorporation of dye-labeled nucleotides into the DNA tag of oligonucleotide-conjugated antibodies, combined with an image-based expression estimation algorithm. Automatic delineation of cell types from multidimensional marker expression and positional data generated by CODEX enabled deep characterization of cellular niches and their dynamics during autoimmune disease both for major and rare cell types populating mouse spleen. A rich source of multivariate data has been generated and provided for the community to further efforts in developing approaches for image analysis, tissue architecture mapping and rare cell type detection.

## Results

### Single base primer extension enables multiplexed antigen staining

DNA provides an ideal substrate for designing molecular tags due to its combinatorial polymer nature. An indexable tagging system whereby tags are iteratively revealed *in situ* by a stepwise enumeration procedure was designed. Antibodies (or other affinity-based probes) are labeled with uniquely designed oligonucleotide duplexes with 5’ overhangs that enable iterative stepwise visualization (**Figure 1A**, **Supplementary Movie 1**, **part 1**). Cells are stained with a mixture of all tagged antibodies at once. At each rendering cycle the cells are exposed to a nucleotide mix that contains one of two non-fluorescent “index” nucleotides and two fluorescent labeling nucleotides. The index nucleotides fills in the first index position across all antibodies bound to the cells. However, the DNA tags are designed such that only the first two antibodies are capable of being labelled with one of the two fluorescent dNTPs - and only if the index nucleotide was previously incorporated. Those two antibodies are then imaged by standard fluorescence microscopy. Then the fluorophores are cleaved and washed away, and the sample is ready for the next cycle where a different indexing nucleotide is used. At the end of the multicycle rendering protocol each pair of antibodies is visualized at a known, pre-defined cycle of the indexing protocol, and the multiparameter image can be reconstructed. The polymerase is paused at the indexing position by omitting one of the indexing (walking) bases from the labeling mix (as done in this study) or potentially by use of reversible terminators (**Supplementary Movie 1, part 2**). Importantly, the system enables multiplexed tissue imaging analysis by means of a standard fluorescence microscope.

**Figure 1.**
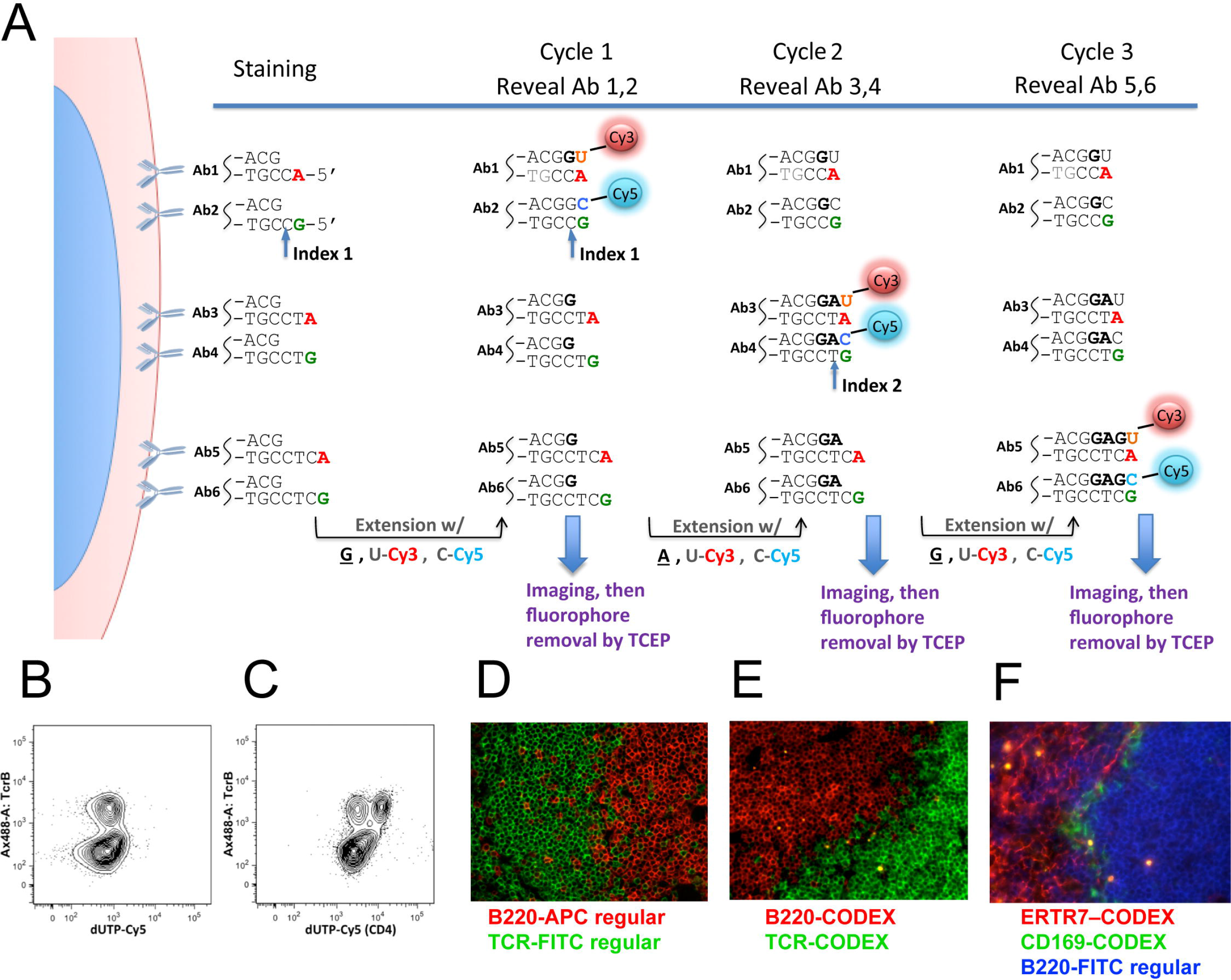
Sequential primer extension on samples stained with DNA barcoded antibodies enables unlimited level of multiplexing. **(A)** CODEX schematic diagram. **(B, C)** Mouse spleen cells were fixed and co-stained with conventional TcRβ Ax488 antibody and CD4 antibody conjugated to CODEX oligonucleotide duplex as in first round of (A). After staining cells were either incubated in extension buffer with dG and dUTP-Cy5 without (B) or with (C) Klenow exo-polymerase. Note that TcRβ-positive T cells in (B) and (C) are indicated by Ax-488 staining. Dependent upon the addition of Klenow, TcRβ-positive CD4 positive T cells are seen as a Cy5 positive subset of TcRβ-positive T cells in (C). **(D)** Spleen cryo-section stained with B cell specific B220-APC (red) and T cell specific TCR-FITC (green) show mutually exclusive staining pattern in the marginal area between B cell follicle and the white pulp. **(E)** Spleen cryo-section stained with CODEX DNA tagged B220 (red) and CODEX DNA tagged TCR-(green) shows staining similar to the one observed with regular antibodies in (D). **(F)** Spleen sections were co-stained with regular B220-FITC and two antibodies (ERTR7 and CD169) tagged with cycell CODEX DNA duplexes. Localization of marginal zone CD169 positive macrophages in the area between the ERTR7 positive splenic conduit of the white pulp and the B220 positive follicular B cells (D) as reported previously has been observed. See also Figure S1, Supplementary Movie 1 Part1

To test the premise of the system, isolated mouse spleen cells were incubated with a CD4 antibody conjugated to an indexing oligonucleotide duplex (as represented by Ab1 in **Figure 1A**). In this trial experiment TCRβ-Alexa 488 was used as a counterstain. A single round of primer extension was done with a mix of unlabeled dGTP and dUTP-ss-Cy5. A cell population positive for both CD4 and TCRβ was observed by flow cytometry. Observation of this population was dependent on the addition of Klenow DNA polymerase to the reaction mixture (**Figure 1B, C**) indicating specific antibody rendering by primer extension. Similarly, in tissue sections, CODEX tag-conjugated antibodies produced lineage-specific staining comparable to regular fluorescent antibodies (compare staining patterns of B220-CODEX and B220-APC in mouse spleen, **Figure 1D-F**).

A simulated multicellular mix was produced by combining 30 aliquots of mouse splenocytes barcoded with pan-leukocytic CD45 antibody labeled with one of 30 distinct CODEX tags (**Figure S1A**). The visualization of the CODEX 15-cycle staining pattern showed even cycle-specific signals, with low background (average signal to noise ^~^ 85:1), efficient (^~^98%) release of fluorophore by inter-cycle TCEP cleavage and no signal carryover between cycles (**Figure S1B-D,F**). Linear regression analysis revealed low signal deterioration (at ^~^0.79% per cycle) and acceptable background (starting at ^~^1.1% and increasing at 0.06% per cycle **Figure S1E,F**).

To further reduce the signal loss associated with accumulation of polymerization errors and to allow larger panels without increasing the length of tagging oligonucleotides, an approach based on primer-dependent subpanels was devised (**Figure S6A**). The feasibility of this design expansion was tested by staining mouse splenocytes with a 22-plex set of antibodies. Each of the antibodies was conjugated to the three versions of the CODEX duplex tag - with same terminated top oligonucleotide and three kinds of the tagging oligo (**Figure S6B**). Thus, every antigen was detected thrice (bringing the overall number of detections to 66) and only after annealing of a panel-specific activator oligonucleotide. We found that the signal for same antibody was consistent across the three primer-dependent batches (**Figure S6C**). Thus panel-activator design extends CODEX to a theoretically unlimited multiplexing capacity, bounded only by the speed and resolution of the imaging process itself and the time required for each imaging cycle.

### Benchmarking CODEX

To validate the quantitative performance of CODEX, cells freshly isolated from mouse spleens were co-analyzed by mass-cytometry (CyTOF) and CODEX using identical 24-antibody panels (**Table S1**). Use of the same antibody clones and the same splenocyte preparation ensured the validity of comparisons. Antigen co-expression signals from CODEX, obtained from image segmentation (see STAR Methods), were consistently similar to CyTOF (**Figure 2B** - for direct comparability, both CODEX and CyTOF data are plotted on a linear scale).

**Figure 2.**
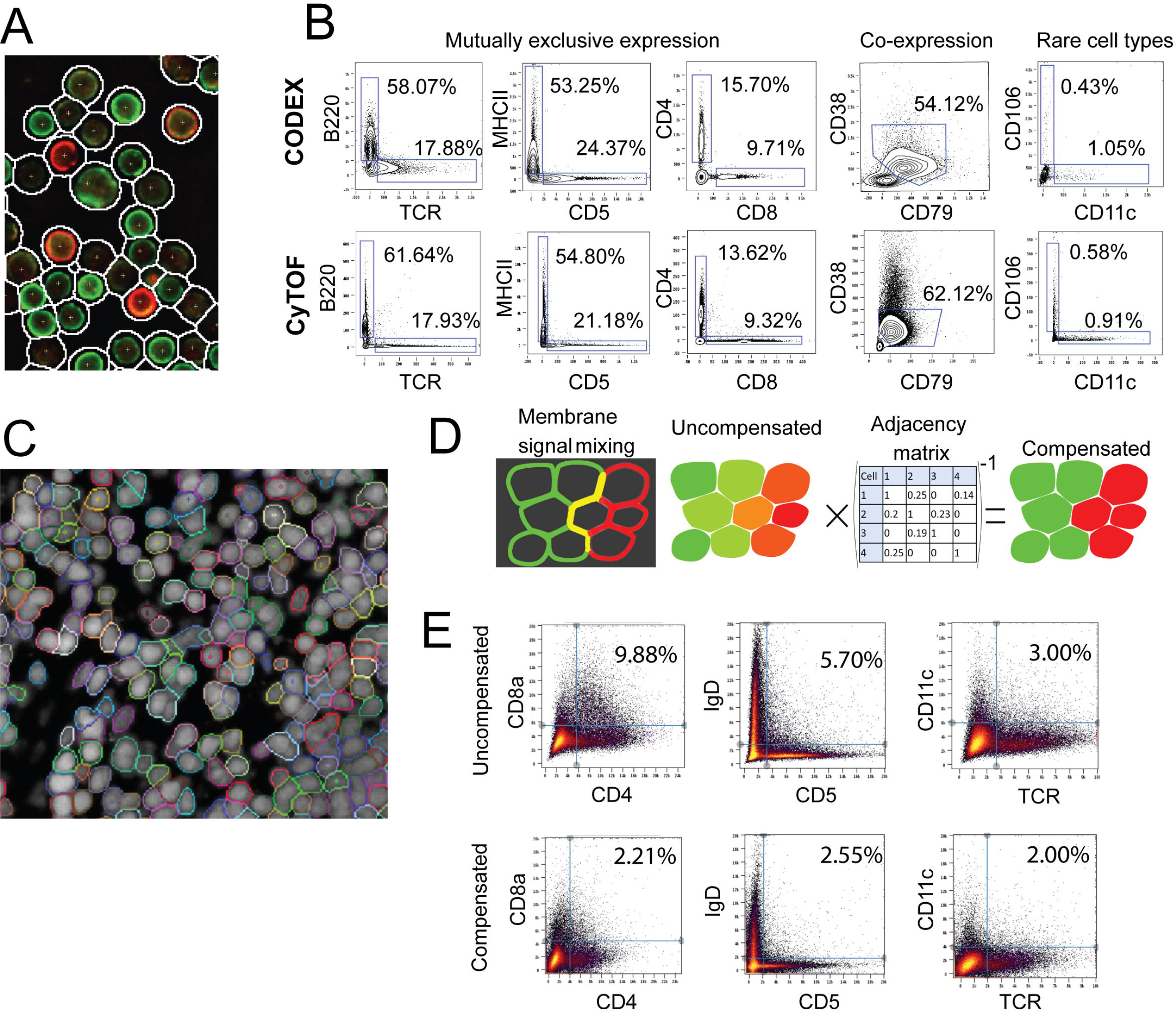
Accuracy of surface marker quantitation by CODEX. **(A)** Microscopic image of mouse splenocytes stained with a 24-color antibody panel, showing one cycle of CODEX antibody rendering. Cell contours show the outlines produced by the cell segmentation algorithm **(B)** Comparison of single-cell expression data derived from dissociated mouse splenocytes on an identical 24-color panel using CODEX and CyTOF. **(C)** Example segmentation in a mouse spleen section based on combining nuclear and membrane (CD45) channel. **(D)** Graphical explanation of the algorithm for compensating the spillover between neighboring cells using a cell-by-cell compensation matrix. **(E)** Biaxial plots of segmented CODEX data acquired in mouse (BALBc) spleen sections. The presence of double positive cells in the upper quadrant is used as an estimate of lateral signal bleeding explained schematically in (D). Three combinations of mutually exclusive lineage markers are shown to demonstrate the range of effect of the compensation algorithm on reduction of lateral signal bleed. See also Figure S2, Figure S3

Consecutively, CODEX was applied to tissue sections. In contrast to dissociated cells spreads, (**Figure 2A**), cells in tissue sections are adjacent to each other—with large fractions of membranes in direct contact (**Figure 2C**). Therefore, neighboring cells can contaminate each other’s signals during the quantification phase (**Figure 2D**). To address this latter challenge, a novel linear algorithm for 3D positional spillover compensation was created. This algorithm is based on the same principles used in fluorescent spillover compensation in traditional flow cytometry, except that our algorithm performs compensation between physically adjacent cell based on their surface contact ratios (**Figure 2**). Indeed, use of this compensation method resulted in a considerable (approximately two-fold) reduction of spillover signal (especially pronounced for CD4/CD8a co-distribution - **Figure 2E**, **Figure 1SL**).

A 30-antibody panel was therefore designed to identify splenic-resident cell types (lymphocytes, macrophages, microvessels, conduit system, splenic stroma; **Figure 3A**, **Table S1**) and applied to the cryo-sections of spleens from wild-type (3 spleens) and MRL/*lpr* mice (6 spleens) (**Figure S4**). Four major classic splenic compartments: red pulp, B-cell follicle, PALS and marginal zone (MZ) (**Figure 3B**) could be easily discerned in CODEX imaging data (**Figure 3A**). A total of 734101 segmented cells were identified, and by means of X-shift clustering (see STAR methods) their expression profiles were grouped into 27 broadly defined phenotypic groups (**Figure 3C-F**, **Table S2**) most of them matching to known cell types. Compared to CyTOF data on splenocytes isolated from non-enzymatically homogenized spleen, CODEX *in situ* analysis produced a similar distribution of cell counts for major cell type. Yet being a non-disruptive technique CODEX identified larger numbers of resident and stromal cell types such as erythroblasts and F4/80 macrophages than CyTOF did (**Figure 3E**). Notably even rare computationally derived cellular phenotypes (e.g. CD4^hi^/CD3^−^/MHCII^hi^ cells and CD11c^+^ B cells) closely matched the cell types previously observed in murine spleen (LTi cells (Robinette et al., 2015) **Figure S3A-C**, **Figure S2C,F,I** and age-associated B cells (ABCs), **Figure S3D,E** and **Figure S2B,E,H**).

**Figure 3.**
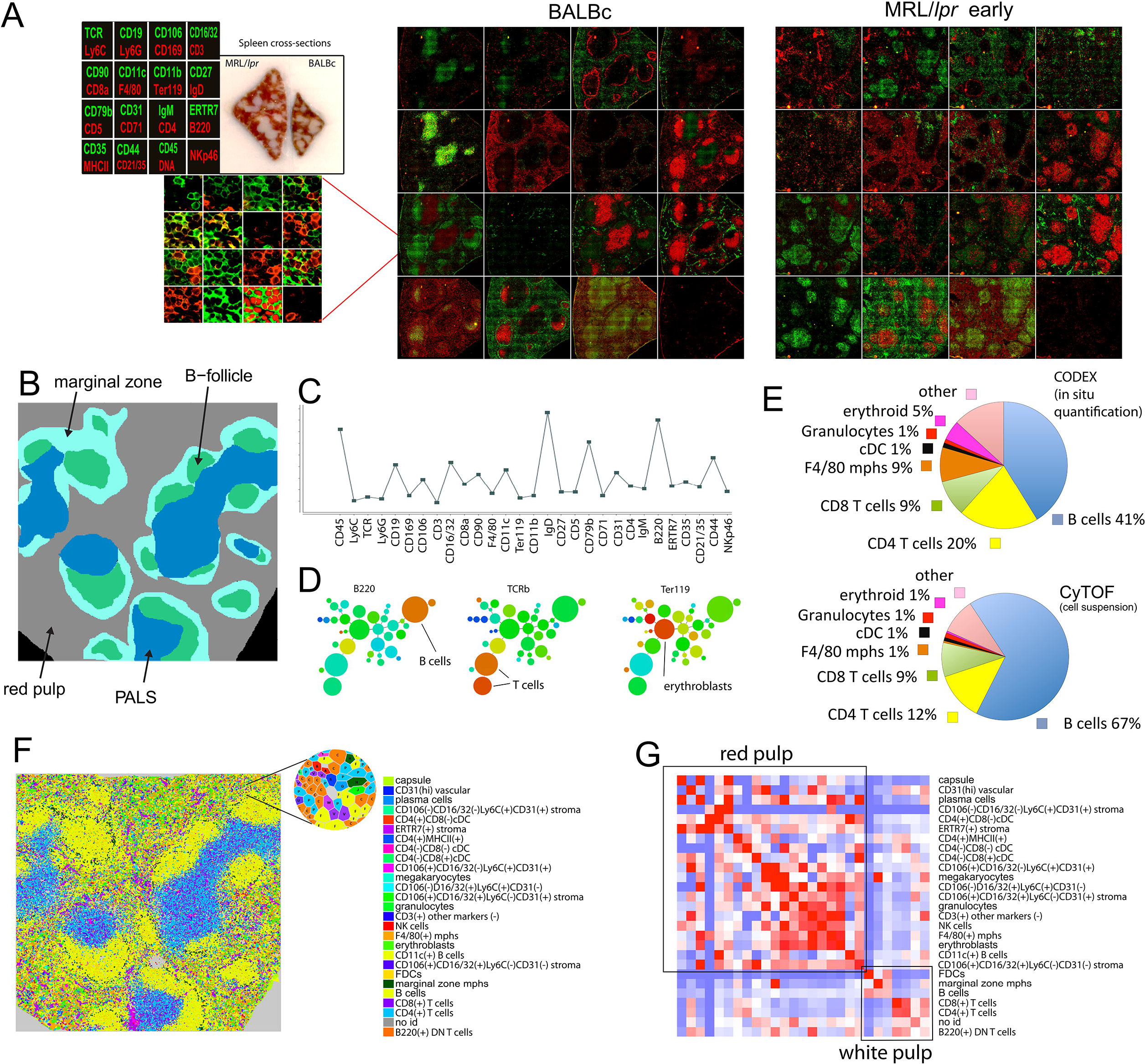
CODEX analysis of mouse spleen cryosections co-stained for 28 antigens. **(A)** Three collated images on the left correspond to: the legend of antibody renderings per cycle; gross morphology photograph of MRL/*lpr* (left) and normal (right) spleen embedded in O.C.T. block prior to sectioning. Green color corresponds to antibodies rendered by extension with dUTP-Cy5, red - dCTP-Cy3 On the right collage of the CODEX multicycle data for normal spleen (BALBc-2) and early MRL/*lpr* spleen (MRL/*lpr*-4). All images are derived from a single scan with a 40x oil objective of an area covered by 63 tiled fields. **(B)** Schematic diagram of major known splenic anatomical subdivisions drawn based on cell distribution in BALBc-1 replicate. **(C)** An exemplary profile of Vortex cluster (B-cells) used for manual matching of clusters to known cell types. **(D)** Minimal spanning tree (MSP) built for all clusters identified by Vortex analysis. On the left middle and right panels the MSPs are colored by expression levels of B220, TCR and CD71 accordingly to indicate location of B-cells T-cells and erythroblasts on the tree. **(E)** Vienne diagram showing for several major cell types their fraction of total cells as identified by CODEX analysis of splenic tissue and CYTOF analysis of isolated BALBc splenocytes **(F)** Post-segmentation derived diagram of identified objects (cells) colored according to cell types in BALBc-1 replicate. Full size diagrams are available for every tissue analyzed in this study are available online (see STAR methods) **(G)** Average cell type to cell type interaction strength heatmap for BALBc samples. Color from blue (<0) to white (around 0) to red (>0) indicates log of odds ratio of interaction (ratio of observed frequency versus expected frequency of interaction). The rows and columns are in the same order (annotation on the right). Black outlines indicate two largely exclusive mega-clusters of cross-interacting cell types loosely matching the cell types populating the red and the white pulp. See also Figure S2, Supplementary Movie 2

### Pairwise and combinatorial (i-niches) statistics of cell to cell contacts in mouse spleen

To provide a high-level view of the cell type interaction landscape, the total counts of contacts between every pair of cell types in the Delaunay neighborhood graph (Gabriel and Sokal, 1969) (**Figure 4A** and **associated Mendeley dataset**) for each condition was determined. The specificity of cell-to-cell interaction was estimated from the “log odds ratio” metric (log-ratio of observed and expected probabilities of contacts between 2 cell types) (**Table S3**). When visualized as heatmaps, this metric revealed a significant non-random distribution of cells in the spleen. In the majority of cases cell types were either selectively associating or avoiding each other (red or blue on the heatmap) pointing to prevalence of specific cell-to-cell interactions in shaping the spleen architecture. The major splenic anatomic compartments were reflected in two large mutually exclusive clusters of positive associations, which appeared to correspond to red pulp and the white pulp, respectively (indicated with black rectangular outlines on **Figure 3G**). For example, a significant positive association was observed between F4/80^+^ macrophages and erythroid cells, as these cell types are both found in the red pulp and are closely associated in so-called erythroblast islands (Socolovsky et al., 2007). An avoidance of interaction was observed between T and B cells, reflecting concentration of these cell types in B cell follicles and PALS, respectively (**Figure 3G**).

**Figure 4.**
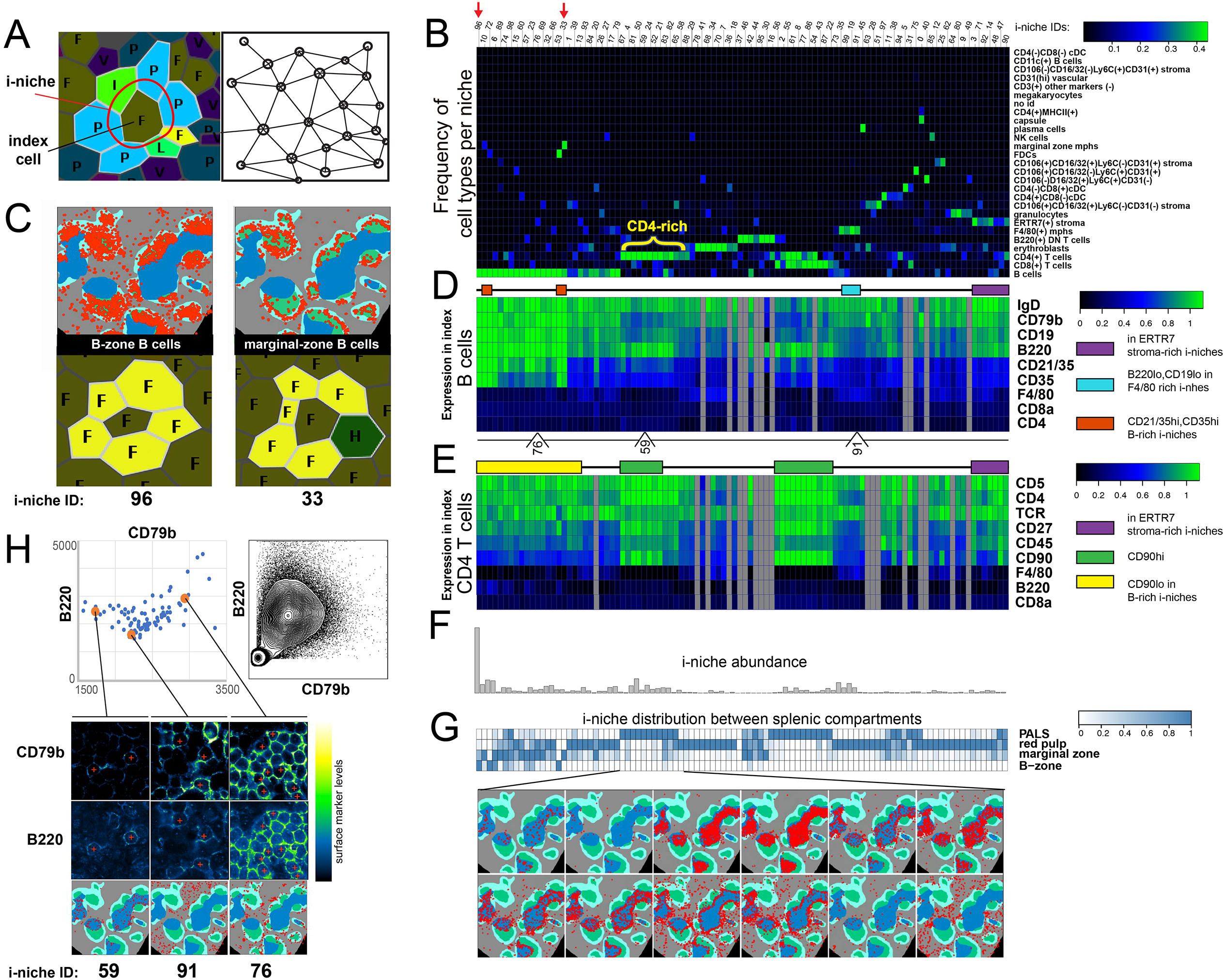
Unbiased identification of i-niches in multidimensional CODEX data. **(A)** On the left - diagram explaining the terminology used for defining i-niche (a ring of first tier neighbors for central cell). On the right - Delaunay triangulation graph used for identification of first tier of neighbors for every cell. **(B)** Heatmap depicting frequency of cell types in 100 types of i-niches identified by K-means (K=100) clustering of all index cells in the dataset (each cell is an index cell for its i-niche) based on frequency of different cell types in the first tier of neighbors. The color indicates the average fraction of corresponding cell type in the the i-niche. **(C)** An example of marginal zone and follicular (B-zone) B cells defined by residence in distinct i-niches (e.g. marginal zone i-niche includes a marginal zone macrophage marked by letter H and green color). Positions of B-cells in each i-niche is marked with red circles over the schematic of BALBc spleen. **(D and E)** Two heatmaps from top to bottom show average expression of selected surface markers measured in a central cell across 100 i-niches (same left to right order as in B) when central cell is B-cells or CD4 T-cell accordingly. The color indicates the relative level of surface marker expression as measured across dataset. Grey columns indicate absence of cells in corresponding niches. Two orange rectangles over top heatmap indicates position of i-niches with high CD35 (containing FDCs and marginal zone macrophages). Cyan rectangle shows location of family of i-niches with high content of F4/80 macrophages and low B220 and CD19 in central B - cell. Purple rectangle indicates family of i-niches enriched with ERTR-7 positive stroma. Below top heatmap location of selected i-niches shown in (E) are indicated. Over bottom heatmap yellow rectangle indicated family of i-niches with dominating presence of B-cells. Two green rectangles indicate family of niches with high levels of CD90 and CD27 in the index CD4 T cells. **(F)** Abundance of 100 i-niches in normal spleen (top bar graph) and **(G)** relative distribution of i-niches between splenic histological subdivisions (PALS, red pulp, marginal zone and B-zone) shown as a heatmap. To illustrate a variety of tissue distribution pattern by i-niches an overlay of selected i-niches over a schematic of normal spleen (BALBc-1) is shown. Heatmap color indicates fraction of corresponding i-niche per splenic anatomic subdivision. **(H)** Top right shows a biaxial plot of flow data for CD79b and B220 measured in isolated splenocytes. Top left shows levels of CD79b and B220 in central B-cells as measured across all 100 i-niches. To illustrate i-niche dependent variability of surface marker expression - images of central cells (marked with red cross) with levels of surface marker indicated in pseudocolor palette are shown for selected exemplary i-niches in the bottom panels. See also Figure S3 K,L

Unexpectedly, a consistently high degree of association was observed between the cells of the same phenotypic class (**Figure 3G**, red diagonal), suggesting that homotypic adhesion constitutes a major force driving the architecture of immune tissue. This observation held true both for the major constituents of white pulp, T and B cells, as well as for rare cell types such as NK cells. Interestingly, even though CD8 and CD4 T cells tended to mix in the PALS, their mutual distribution was nonrandom and consisted of intertwined threads of homotypic cells (**Figure S3H**). Interestingly, as an aside, similar structures could be reproduced in vitro by incubating heterotypic mixtures of sorted splenic cell populations (**Figure S3 I,J**). These data suggest that homotypic cell association might be an important driver of the white pulp substructure and is worth further investigation.

The precision *in situ* cytometry analysis of CODEX data allowed enumeration of cellular contexts in a manner not possible previously. We define here an indexed “niche” (i-niche) as a ring of cells (excluding the central, or here defined as “index” cell) in no specific circumferential order that are Delaunay neighbors of the index cell (**Figure 4A** and STAR methods). We identified 100 of the major i-niches (by K-means clustering) according to the relative frequency of the identified cell types present in the ring of cells surrounding the index cell (**Figure 4A,B** and **F**). Most i-niches could be readily mapped into one of major anatomical compartments of the spleen (B cell follicle, PALS, marginal zone, or red pulp - per **Figure 4G**). In most cases, any given i-niche resided within a single anatomical compartment (although several i-niches were observed in more than one compartment), and every splenic compartment was populated by many i-niches (**Figure 4G**).

In our definition the index cell in the center can be ***any*** cell. Thus i-niches enabled subsetting the common cell types based on cellular context (microenvironment). For example, B-cells surrounded by only B-cells (red arrow **Figure 4B**, i-niche #96) can be seen primarily in the follicular zone B cell region (**Figure 4C**, **left panel**) while presence of CD169 positive marginal zone macrophages mapped the B cells in such i-niches to the marginal zone (red arrow **Figure 4B**, i-niche #33, **Figure 4C**, **right panel**). In the case of T cells, CODEX data enabled precise selection of an important migratory subset of T-cells known to be residing in ERTR7 enriched niches (Burrell et al., 2015) (in **Figure 4B,E** see the purple rectangle indicating a family of niches where index T cells contact ERTR7 stroma; as well as **Figure S5A,B**). Taken together, we see that surface marker expression alone is insufficient to associate many cell subsets with a given tissue subcompartment (e.g. CD4^+^ T cells can be found both in the PALS and in the red pulp). However i-niche designation does provide such mapping data (most of i-niches were enriched within a specific splenic subdivision **Figure 4G**) and as such T cells associated with ERTR7 positive stroma in fact localize primarily to the red pulp (**Figure S5A,B**). This raises interesting questions—can new cell types, or functional subsets, be discerned by this approach? What is the frequency of a repeated i-niche structure that must be observed to suggest a function? And what would constitute a proof that a given i-niche corresponds to a new cell type or functionality?

### Cell surface marker expression depends on local neighborhood

One approach to address these latter questions is to consider the phenotypes of the index cell in various i-niches. We observed that for several index cell types (e.g. for B and T cells) there was significant biasing of the surface marker expression depending on the i-niche in which the index cell resides (e.g. see selected cases marked with colored rectangles above the heatmaps in **Figure 4D** and **E**). To assure that it was not a quantitation artifact we mapped back to the image the index B-cells where the levels of CD79b (a co-activator chain of the B cell receptor complex) and B220 (a splice isoform of CD45 membrane phosphatase) would be niche-dependent. We found that the index B cells that were B220^int^, CD79b^lo^(i-niche “59”) resided on the boundary between the PALS and the follicles (**Figure 4H** image montage on the bottom). Index B cells that were B220^int/hi^, CD79b^int^ (i-niche “91”) were mostly found in the red pulp. And, B cells that were B220^int/hi^, CD79b^hi^ (i-niche “76“) were yet different again and were found at the boundary of the red pulp and the follicles. When measured in cell suspensions, CD79b is co-expressed with B220 at various ratios (see CyTOF plot single cell splenocytes **Figure 4H**, top right panel). Such a distribution of expression is sometimes attributed to staining variability, measurement noise, or a simple lack of understanding of the underlying biology. However, as seen on **Figure 4H**, upper left panel, there is a non-random pattern of CD79b and B220 expression across the central cell of the corresponding i-niches and, depending on the B220/CD79b levels, the i-niches (the central cells) map to specific regions in the splenic architecture (**Figure 4H**, lower panel). These observations suggest that the spread of the CD79b-B220 levels as well as of other marker levels on splenic B-cells could be, to a large degree, accounted for by the niche composition around those B-cells - and that the expression levels on these cells might be influenced by (or influences) the cells in their immediate surrounding.

The overall utility of the i-niche in determining any given surface marker expression value for an index cell was evaluated by constructing a linear regression model of marker expression using both the cell type identity and the i-niche constituency in a two-featured variable model (the other variable being the cell type identity). Notably, adding the i-niche information as a dependent variable significantly improved the fitness of the model for all markers (**Table S4**) with highest improvement F-values for CD90, B220, CD21/35, and ERTR7 and the lowest prediction rates for Ly6G, CD5, CD11b, CD5, and TCRβ. Thus, the high variability in B220 expression levels are highly related to the i-niche in which the B cell resides. In other words, B220 expression levels can be location-specific, and are dependent on the i-niche partners. As a counter point, the data also shows that i-niche does not reliably predict expression of other proteins, such as CD5 or TCRβ, the expression levels of these receptors is relatively constant across the i-niches (**Figure 4E**). This result quantitatively demonstrates that the i-niche (neighbors) determines a significant proportion of variance in the expression of certain markers. Overall, we observed that that many splenic cell types populate a wide variety of i-niches (**Figure S3K,L**), suggestive of a multiplicity of functional state for any given immune cell type. Further, tissue locale (i-inches) is a powerful indicator of potential differential function (to the extent tissue locale drives function) and these deterministic changes in surface marker protein expression are surrogate indicators of this locale or function.

### Changes in splenic composition associated with disease progression

A comparable region of spleen was visualized by CODEX for 3 normal BALBc spleens, and 6 spleens from MRL/*lpr* mice. Image segmentation revealed strong variation in cell counts between the norm and the disease (**Figure 5B**) for most (19 out of 27) of the cell types identified by X-shift clustering. Examples include a dramatic increase in CD71^+^ erythroblasts (green cells on **Figure 5A** maps), a reduction in numbers of B cells and FDC, and increases in so-called B220^+^DN T cells (CD4/CD8 double-negative B220^+^ T cells), which have been previously characterized as a hallmark of the MRL//pr progression (Koh et al., 1995) which could also be identified by FACS (**Figure S1F,G**), thus ruling out the possibility that this unusual cell type being a result of image segmentation errors. These and other changes were used to broadly classify the MRL/*lpr* spleens into early, intermediate, and late disease stages (**Figure S4**).

**Figure 5.**
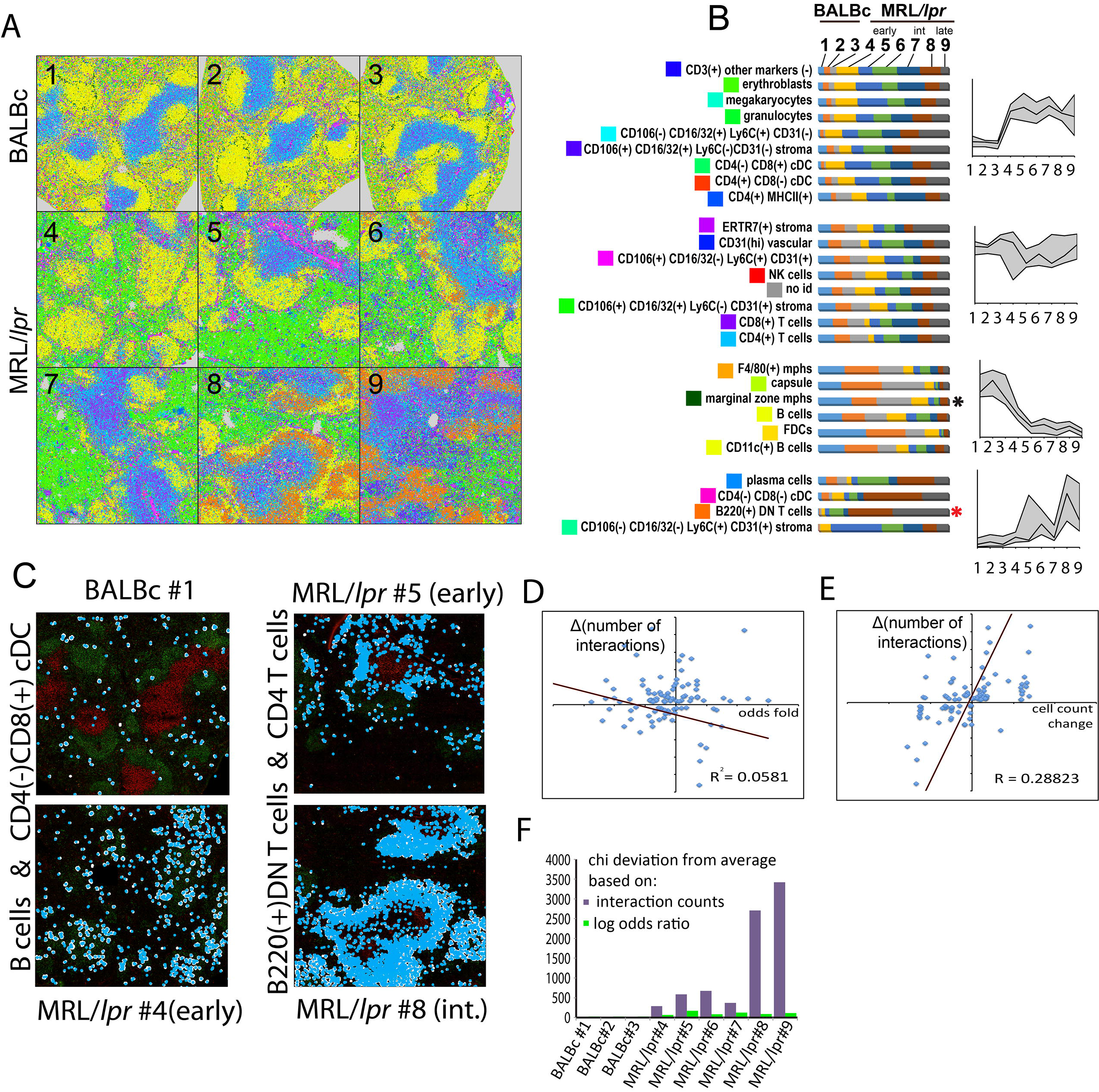
Autoimmune disease drives changes in splenic composition and cell-to-cell interactions. **(A)** Post-segmentation diagrams of all objects (cells) colored according to cell types (see color map in Figure 3F) for all normal and MRL/*lpr* tissue sections imaged in the study. Full size diagrams are available for every tissue analyzed in this study are available online (see STAR methods) **(B)** Stacked bar graphs show dynamics of cell counts across dataset for manually annotated Vortex clusters (cell types - on the left) across progression from normal to afflicted spleen. Colored bar sections indicate fraction of the total cells as detected at a particular stage/samples (1-9 annotation on the top). Cell types were split into four types according to the dynamics of counts across dataset as represented by average relative (normalized to 1) count - see line graphs on the right, x axis corresponds to stage/sample id. **(C)** Two examples of change in cell-to-cell interaction frequency during disease progression - between the B cells and dendritic cells in normal and early MRL/*lpr* spleen and between B220+ DN T cells and CD4 T cells during progression from early MRL/*lpr* to intermediate. **(D)** Co-distribution of odds ratio log-fold [log(odds ratio in early MRL*/lpr*)- log(odds ratio in BALBc)] on X axis and change in counts of interactions for early MRL/*lpr* versus control (BALBc) comparisons (on Y axis). **(E)** Co-distribution of cumulative cell frequency change [celltypel freq. change + celltype2 freq. change] on X axis and change in counts of interactions for early MRL/lpr versus control (BALBc) comparisons (on Y axis). **(F)** Bar graph showing Chi square values across conditions computed for odds ratio and direct interaction counts. See also Supplementary Movie 2, Figure S5

Despite consistent presence of “homotypic interactions” diagonal and larger cell-adjacency clusters corresponding to red and white pulp in odds ratio heatmaps across disease (**Figure S4**), a deeper statistical analysis revealed many disease-associated changes in frequency of contacts between cell types (see **Table S3**). Among the changes we observed an increase in interaction between B cells and CD4^−^/CD8^+^ cDC in the early MRL/*lpr* spleen compared to normal, (**Figure 5C** left panel) suggesting an increase in B-cell activation. We also observed a higher interaction frequency of granulocytes with T cells (**Figure S5C**), dendritic cells (**Figure S5D**) and erythroblasts (**Figure S5E**), and a higher number of contacts between erythroblasts and various kinds of stromal cells, as well as B220^+^ DN T cells (**Table S3, Figure S5F,G**). In the intermediate and late stage MRL//pr spleens, there was a significant increase in interaction of B220^+^DN T cells with CD4^+^ T cells (**Figure 5C** right panel), CD8^+^ T cells, erythroblasts, and a variety of other cell types compared with numbers of these interactions in the early MRL/*lpr* stage (**Table S3** and **Figure S5G-I**, see online resource, STAR methods, for more examples). So, while there was no obvious gross rearrangement of the tissues, many homotypic and heterotypic cell-cell associations were altered, prompting a key question: what are the main factors driving this disruption?

### Disease driven change in cell counts determines the frequency of specific cell-to-cell contacts

What could be the drivers of changes in frequency of pairwise cell-cell contacts? If the kinetics of pairwise cell contacts, follows a rate law — one possibility would be that modulation of specific cell-to-cell interaction potential—or “attraction” (for which the odds ratio score was used as an estimate across this study) is the main driver. In other words, it would be expected that when the affinity of such an interaction goes up, the fraction of interacting cells of a given cell pair would increase. At the same time, even in the absence of change in cell-to-cell affinity, the absolute number of the cell-cell pairs (defined here as cell pair aggregates, or CPAs) and the number of interacting cell pairs should correlate with the frequencies of interacting cell types (analogous to concentrations in the rate law equation). Importantly, the latter scenario could be as biologically significant as the former. Finally, some of the cell type contacts may be observed due to low cellular motility of randomly meeting cells. Such interactions would not produce spatially defined sub-splenic CPAs and would have and odds ratio close to 1.

The perturbation introduced to normal splenic composition with MRL/*lpr* genotype allowed us to examine the mechanisms implicated in transition from normal to diseased spleen. In short, we found that, for most cell-cell pairs observed, the mutual attraction (as quantified by the interaction log odds ratio), was not the primary determinant driving the change in counts of interacting cell pairs between the MRL/*lpr* and the norm. In **Figure 5D** we plot the change in counts of interactions of two cell types (e.g. A:B) between the MRL/*lpr* and the normal BALBc spleens. Each dot represents a pair of cell types. The value on the Y axis is the difference in the total number of observed interactions between BALBc and MRL/*lpr*. The X axis shows the difference between log odds ratios of interactions between the same conditions. There was no overall correlation observed (R^2^ = 0.058). In contrast, we observed a correlation (R^2^ = 0.288) between the cell count changes and the interaction changes (**Figure 5E**). In agreement with those observations, we saw that out of the 26 top scoring (FDR < 0.05 and change in absolute interaction counts > 150) cell type pairs of this cross comparison only 2 showed corresponding significant (FDR < 0.05) change in odds ratio score. Curiously these two interactions with a modest 1.5 times increase in interaction count and, concomitantly, a ^~^0.8 increase in log odds ratio score were the ones between the CD4 or CD8 T cells and ERTR7^+^ stroma (see **Table S3**, rows 6 and 7, and **Figure S5A,B**). Visually they appeared as persistent co-clustering of T cells with ERTR7^+^ stroma despite the overall drop of T cell numbers in the “early” MRL/*lpr* samples. Curiously, ERTR7 positive fibers of splenic stroma as well as ERTR7 protein itself were recently shown to be critically involved in T cell trafficking (Burrell et al., 2015), suggesting that this increase in the spatial association could be reflective of the T cell activation.

For the rest 24 of the 26 changing interactions mentioned above at least one of the cells of the pair was scored as significantly (FDR<0.05) changing the frequency across scored conditions (**Table S3** last column of the “EarlyMRL vs BALBc control” spreadsheet). We therefore conclude that—at least in the diseased state of early stage MRL/*lpr*—most of the change in counts of cell-cell interactions are driven simply by increases or decreases in cell type frequencies.

As an additional evidence, χ^2^ statistics were used to compare the total magnitude of changes in pairwise cell type interaction matrices (total interaction count) versus changes in log-odds ratio matrices (propensity for non-random interaction). The χ^2^ deviation (sum of squares of z-score-normalized values) was computed for each disease matrix compared to the control. In every case, the χ^2^ values of cell interaction matrices were larger than of the respective log odds ratio matrices of the same biological sample (**Figure 5F**). This suggests that as the cell type frequencies change due to disease progression, the absolute numbers of interactions change dramatically whereas the frequency-normalized likelihoods of cell interactions change to a much smaller extent indicating a great degree of robustness of the ‘design principles’ of the splenic tissue and that many of the more dramatic disease-associated variations occur primarily through the shift in cell numbers.

This analysis implies that the degeneration of the tissue integrity in the MRL disease largely follows dramatic changes in cell type frequencies. At the same time, there were notable exceptions from this trend, where the changes in observed cell type pairing frequencies could be largely explained by shifts in the cell type interaction likelihoods (log-odds ratios). While further work is required to determine which of these changes are instrumental to the MRL disease state, our findings suggest that such differential analysis can be applicable in other diseases, and possibly, could be used to discover cell type interactions that are targetable from a therapeutic standpoint.

### Reorganization of cells in disease-associated tissue substructures

We catalogued the cell-cell interaction “connectivity” in a circular correlation diagram. Rarely, if ever, there was any cell type found adjacent to only one other type of cell. The highest degree of connectivity was observed for the most abundant cell types such as B cells in normal spleen and erythroblasts (**Figure 6A**) in early MRL/*lpr*. This high connectivity in turn led to large effect on i-niches caused by changes in cell numbers associated with progression of disease from normal to autoimmunity. Most dramatic changes in cell frequencies were the increase in erythroblasts in the early MRL/*lpr* and the emergence of B220^+^ DN T cells in late MRL/*lpr* - which were associated with appearance of novel i-niches relative to the normal spleen (spatial localizations of B220^+^DN T cell dominated i-niche 18, erythroblast driven i-niche 29 and B-cells rich i-niche 96 are shown on **Figure 6B** and their cell type composition is shown on heatmap on **Figure 6C**). A corollary to this is the question of whether the presence of these cells, and new i-niches dependent on these cells, somehow changed the observable biology of the cells they contact? We found some examples supporting that, whereby the proximity of CD4 T cells to B220^+^DN T leads to CD4 T cell activation in spleens of MRL/*lpr* mice: **Figure 6C** shows increased levels of CD27 expression in CD4 T cells present in i-niches dominated by B220^+^ DN T cells (**Figure 6C** red circle).

**Figure 6.**
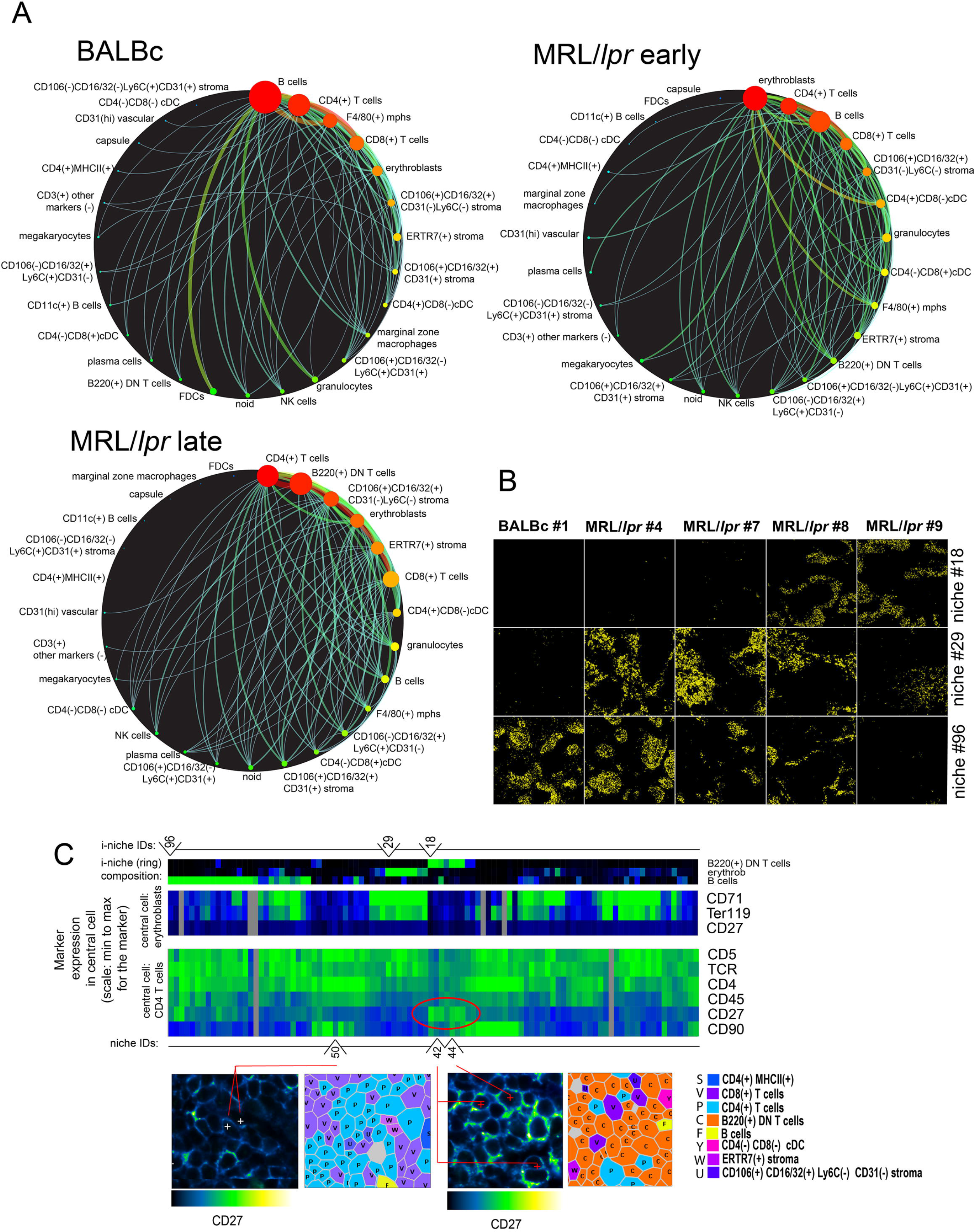
Differential effect of disease over i-niche presence across dataset. **(A)** Cell interaction networks built for BALBc early MRL/*lpr* and late MRL/*lpr* based on number of contacts observed between two cell types (only connections with more then 150 interactions per sample are shown on the diagrams). Thickness of connection correlates with number of contacts size of the node indicates number of cells per condition. **(B)** Evolution of i-niche abundance across dataset. Selected three i-niches (marked above heatmap in (C) depicting i-niche composition) differentially represented across dataset (changing between norm and disease) are shown. Yellow circles overlaid over blank rectangle corresponding to imaged area indicate location of i-niche. **(C)** Top heatmap shows frequencies of B220^+^ DN T cells, erythroblasts and B-cells in the i-niche rings. Line above top heatmap indicates the composition of i-niches 18, 29, and 96 described in (B). Color scheme is the same Middle heatmap indicates expression of selected markers when the i-niche central cell is an erythroblast - primarily to show that CD27 is not expressed on erythroblasts in the vicinity of B220^+^ DN T cells. Bottom heatmap indicates expression of selected markers when the i-niche central cell is a CD4 T cell. The color schemes in these three heatmaps is the same as in heatmaps on Figure 4B,D,E. Red oval outline pinpoints i-niches with elevated CD27. Note that these i-niches as indicated by top heatmap have B220^+^ DN T cells as a prevailing component. Lower panels show examples of central cells in i-niches marked under the lower heatmap. i-niche 50 is an example of i-niche without B220^+^ DN T cells. Central cell does not express high CD27. i-niches 42 and 44 have high frequency of B220^+^ DN T cells and accordingly central cells express high CD27. See also Figure S3 K,L

Other cell types noticeably changed their characteristic distribution and their propensity to engage, or evade, specific cell-to-cell contacts (as estimated by odds ratio score) during disease progression. For example, cells of CD106^+^CD16/32^−^Ly6C^+^CD31^+^ phenotype were randomly distributed in the red pulp of normal spleens, but were found to aggregate in the areas proximal to the marginal zone of the MRL/*lpr* white pulp (**Figure S2D,G,J**). This re-distribution correlated with erythroid proliferation and reduced odds ratio score for the interaction of CD106^+^CD16/32^−^Ly6C^+^CD31^+^ and erythroblasts in lupus spleens (**Table S3**).

### Automatic definition of disease-associated areas in tissue architecture

As noted, the analysis reveals that the development of the autoimmune disease in mice (as exemplified by MRL/*lpr* lupus) is associated with vast rearrangement of normal spleen architecture, which is likely to cause loss of cell-cell contexts normally hosting the cells crucial for proper splenic function, as well as the observed emergence of novel i-niches that are not found in the normal BALBc spleen. Additionally, certain i-niches were sequestered to specific anatomic compartments of the spleen, which allowed us to use such i-niches as reference points to quantitatively monitor high-order morphological changes. The i-niches that in normal spleen were localized to one distinct compartment (more than 90% of central cells reside within a particular splenic compartment) were used to evaluate the dynamics of splenic cells associated with progression of autoimmune disease (**Figure 7A**, **middle heatmap**). This analysis confirmed the dissipation of the marginal zone starting from early stages of MRL/*lpr* and revealed a progressive distortion of PALS. Curiously, depending on whether a i-niche was based on F4/80 macrophages or primarily contained erythroblasts, the red pulp appeared to reorganize in the diseased tissue (**Figure 7A**, **right heatmap**), pointing to the fact that more than one compartment-specific niche is required to reliably trace the fate of specific anatomic compartments. In many cases the definition of subsets/morphological units constituting the tissue is subjective, yet this study employed niches that were algorithmically defined. Therefore, using niches as markers of morphology can quantitatively monitor the changes of high-order anatomic architecture.

**Figure 7.**
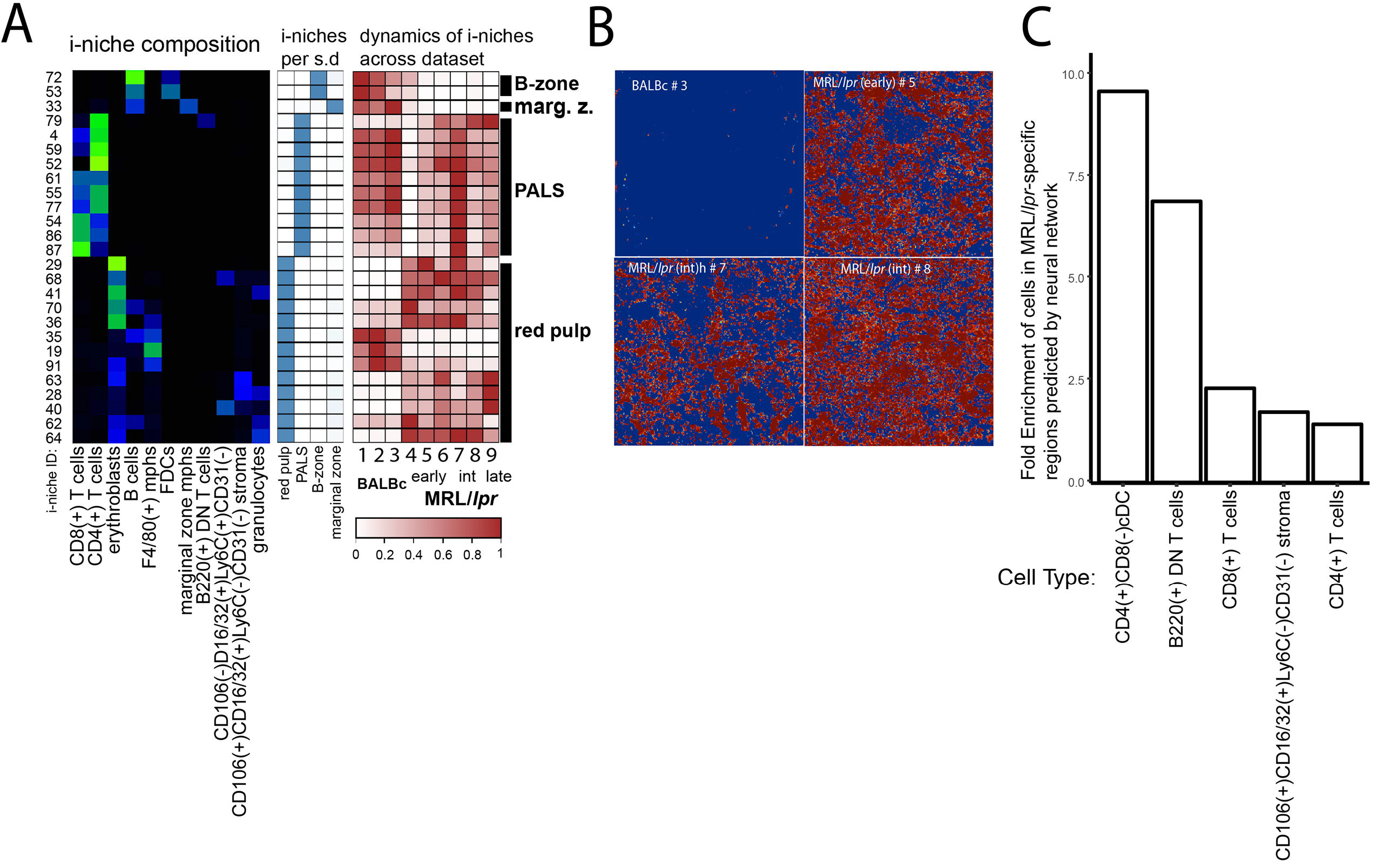
i-niches and neural nets provide unbiased way for disease monitoring. **(A)** Selected i-niches (green heatmap shows i-niche composition, color scheme same as in Figure 4B) were chosen based on high (>90%) presence per single histological subdivision (blue heatmap color scheme same as in Figure 4G). Abundance of these i-niches (brown heatmap, color indicates relative abundance of corresponding i-niche as measured across full dataset) was used to judge the preservation or decay of a histological splenic subdivision corresponding to selected i-niches. **(B)** Red color over blue rectangle indicates regions of interest (MRL/*lpr*-specific regions) predicted by neural network in entire spleen images. From top left, clockwise: BALBc #3, MRL/*lpr* #5, MRL/*lpr* #7, MRL/*lpr* #8. **(C)** Cell types enriched (FDR < 0.1) in MRL/*lpr*-specific regions (in red in B) predicted by neural network. See also STAR Methods “Neural network training“

To automatically isolate the specific local combinations of expression patterns characteristic of the disease state, a fully convolutional neural network was trained to distinguish image patches from normal and MRL/*lpr* mice. The neural network operated by identifying, in each training image patch, the specific areas that corresponded to the disease state.

The neural network highlighted the regions in each multiparameter spleen image that corresponded to the disease state (**Figure 7B**), despite having seen no images from these spleens during training. To investigate the specific features learned by the neural network, the cell-type compositions of the regions identified as diseased versus those regions identified as normal were compared. There was significant enrichment of several cell types in these regions (**Figure 7C**). Although some cell types enriched in diseased regions, for example B220^+^ DN T cells, were present only in the diseased tissue, the most highly enriched cell type (CD4^+^/CD8^−^ cDCs) were present in both the disease state and the healthy state.

To assess the specific contextual changes recognized by the neural network, the local neighborhoods of the CD4^+^/CD8^−^ cDCs that the neural network found to be enriched in MRL*lpr* regions were analyzed. In these neighborhoods we observed a significant enrichment of other CD4^+^/CD8^−^ cDCs, as well as significant depletion of CD106^+^/CD16/32^+^/Ly6C/CD3r stromal cells (FDR < 10“^−7^). This suggests that the neural network had identified an altered context for CD4^+^/CD8^−^ cDCs (distant from stromal regions) as a key descriptor for the disease. Thus, the neural network approach described here enabled both automatic classification of samples according to disease state and an automatic identification of high-dimensional regions of interest and corresponding cellular niches.

## Discussion

Here the feasibility of polymerase-driven highly multiplexed visualization of antibody binding events to dissociated single cells as well as tissue sections (CODEX) was demonstrated and benchmarked. Critically, CODEX enables co-staining of all antigens simultaneously with the staining iteratively revealed by primer extension cycles wherein no diminution of epitope signal detection was observed. A consistent performance of CODEX in co-detecting up to 66 antigens was demonstrated, and the “activation primer” - based extension of the system could enable a potentially vast expansion of CODEX multiplexing capacity. For the current method fresh-frozen tissue was used yet at a cost of testing an extensively broader collection of clones we have recently succeeded in adapting the procedure to FFPE archival tissue (manuscript in preparation). We believe this will open the large retrospective collection of FFPE samples from clinical cohorts to multidimensional cytometric analysis.

The CODEX platform can be performed on any three-color fluorescence microscope enabling conversion of regular fluorescence microscope into a tool for multidimensional tissue rendering and cell cytometry. CODEX completes a 30-antibody visualization in approximately 3.5 hours. Modifications to the technology that increase the measurements per cycle, reduce the cycling time, faster imaging methods such as light sheet microscopy, or an increased size of the imaging the field of view offer potential opportunities for increasing the depth and speed of the visualization process. Given the low cost of converting a scope to this platform (enabled by a simple fluidics device for automated sample washes in a customized microscope stage) CODEX technique could enable deep studies of various tissue models even with limited resources and instrumentation.

The unique set of algorithms described here successfully identified individual cells in the crowded environment of lymphoid tissue by relying both on the information from nuclear and the membrane staining. An accurate quantification of single-cell expression data was obtained directly from the images by creating a special algorithm for positional spill compensation. As of today, this algorithm is only applicable to uniformly distributed surface markers. Future changes might be required to accommodate markers that follow a different distribution, i.e. localized to lipid rafts or immune synapses. Nevertheless, the use of this algorithm enabled us to extract FACS-like data from tissue imaging and leveraged the automated phenotype mapping framework previously developed for CyTOF and multicolor FACS.

Performance of CODEX on tissue sections was validated in analysis of spleen sections of normal and lupus afflicted mice (MRL/*lpr*). Much like with conventional flow cytometry, CODEX discerned all major cell types commonly observed in mouse spleen. Moreover, application of X-shift phenotype-mapping algorithm (Samusik et al., 2016) tailored to parsing the multidimensional single-cell data enabled the detection of rare cells types (such as CD4^hi^ MHCII^hi^ (Lti) cells, CDllc(+) B cells (ABC cells)) and simultaneously placement of those cells in the tissue architecture. Cell interaction analysis with CODEX recapitulated known features of splenic tissue architecture and revealed that most splenic cell types were frequently in homotypic interactions—which might underscore a novel driving principle of lymphoid tissue architecture. Further, an important principle was derived from the data-driven i-niche analysis, i.e. we have established that certain markers, such as B220, CD79b or CD27 exhibited significant changes in expression levels depending on the tissue context in which the cells reside. As clearly observed in experiments that drove **Figures 4** and **6**, cell populations that would otherwise be thought of as ‘broadly’-expressing a given marker set (**Figure 4H**), in fact were composed of multiple subphenotypes that correlated with the i-niche identity. In other words, what immunologists previously thought of as a single cell type could be subdivided into more subtle cell subsets that are defined by the neighborhood in which they reside. We leave open the question of whether the cells with different properties are attracted to a set of neighbor cells, or a given expression level of markers attracts the neighbors, or some dialectics thereof. What is clear, however, is that there are more subtle phenotypes in tissues than previously assumed, and that future developments future of the technology and the algorithms will shed more light on these phenomena.

CODEX enabled a quantitative description of autoimmune-related changes in the splenic tissue architecture. Among hallmarks of MRL/*lpr* progression were dissipation of marginal zone, disintegration of PALS, invasion of red pulp with erythroblasts and the infiltration of mixed-identity B220^+^ DN T cells, which, interestingly, localize in a niche in between PALS and the B cell zone and in the marginal zone. A contact-dependent effect of B220^+^ DN T cell on CD4 T cells reflected in increased levels of activation marker CD27 was observed. An account of statistically significant differences in frequency and strength of pairwise cell type contacts was created. From these observations and their quantitative analysis, we concluded that it is largely the change in cell numbers rather than in cellular interaction strength (estimated from ratio of observed to expected probability of interaction) that is involved in reorganization of spleen during transition from norm to autoimmunity. We show how iniche statistics can be used to account for the list of disease driven changes in sub-splenic anatomy. We also show that disease-associated areas of the tissue can be identified independently of the image segmentation, by applying a convolutional neural network to the multidimensional image data, even after training on just one sample.

Recent advances in genomics suggest that despite the vastness of a genetic repertoire—there exist only a limited number of cellular states with a concomitantly limited gene expression pattern. These countable, limited, patterns are reflected in expression of surface marker phenotypes recognizable as cell types. It is therefore reasonable to suggest that cell-to-cell interactions should be limited as well and falling into repeated patterns. By this token, the data collected in this study lays the foundation for a pan-cellular reference database defining cellular types not only by identities of proteins expressed but also by definitions for specific cell-to-cell interactions. We performed deep characterization here for normal and diseased tissue from such a perspective of cell-cell arrangements and present here, for the research community, a large (^~^700,000 cells) public dataset encompassing segmentation, quantification, and, most uniquely, spatial data from normal and disease-afflicted spleens (http://welikesharingdata.blob.core.windows.net/forshare/index.html). Further analysis of data could enable advances in understanding of clinically relevant cell interactions in immune tissues as well as development of computational algorithms for tissue cytometry and digital pathology.

## Acknowledgements

We thank A. Trejo and A. Jager for technical assistance. We are grateful to Peter Jackson for advice and access to the microscope used for this work. This work was further supported by grants to GPN: U19 AI057229, 1U19AI100627, Department of Defense (CDMRP), Northrop-Grumman Corporation, R01CA184968, 1R33CA183654-01, R33CA183692, 1R01GM10983601, 1R21CA183660, 1R01NS08953301, OPP1113682, 5UH2AR067676, 1R01CA19665701, R01HL120724, R01HL128173 (NIH subaward under VCU); Stanford Center Cancer Systems Biology grant U54-CA209971; GPN is supported by the Rachford & Carlotta A. Harris Endowed Chair.

## Author Contributions

Conceptualization: Y.G., N.S. and G.P.N.; Methodology: Y.G., N.S. and J.K.-D.; Software: N.S. and S.Bh.; Validation: Y.G., N.S., J.K.-D. and G.V. ; Formal Analysis: Y.G. and N.S.; Investigation: Y.G. and N.S.; Resources: Y.G., N.S., G.V., S.B., and M.H.; Data Curation: Y.G. and N.S.; Writing Original Draft: Y.G. and N.S.; Writing Review and Editing: Y.G. and N.S. and G.P.N. ; Visualization: Y.G. and N.S.; Supervision: Y.G. and G.P.N.; Project Administration: Y.G. and G.P.N.; Funding Acquisition: G.P.N.

## Supplementary Figures and Legends

**Figure S1.**
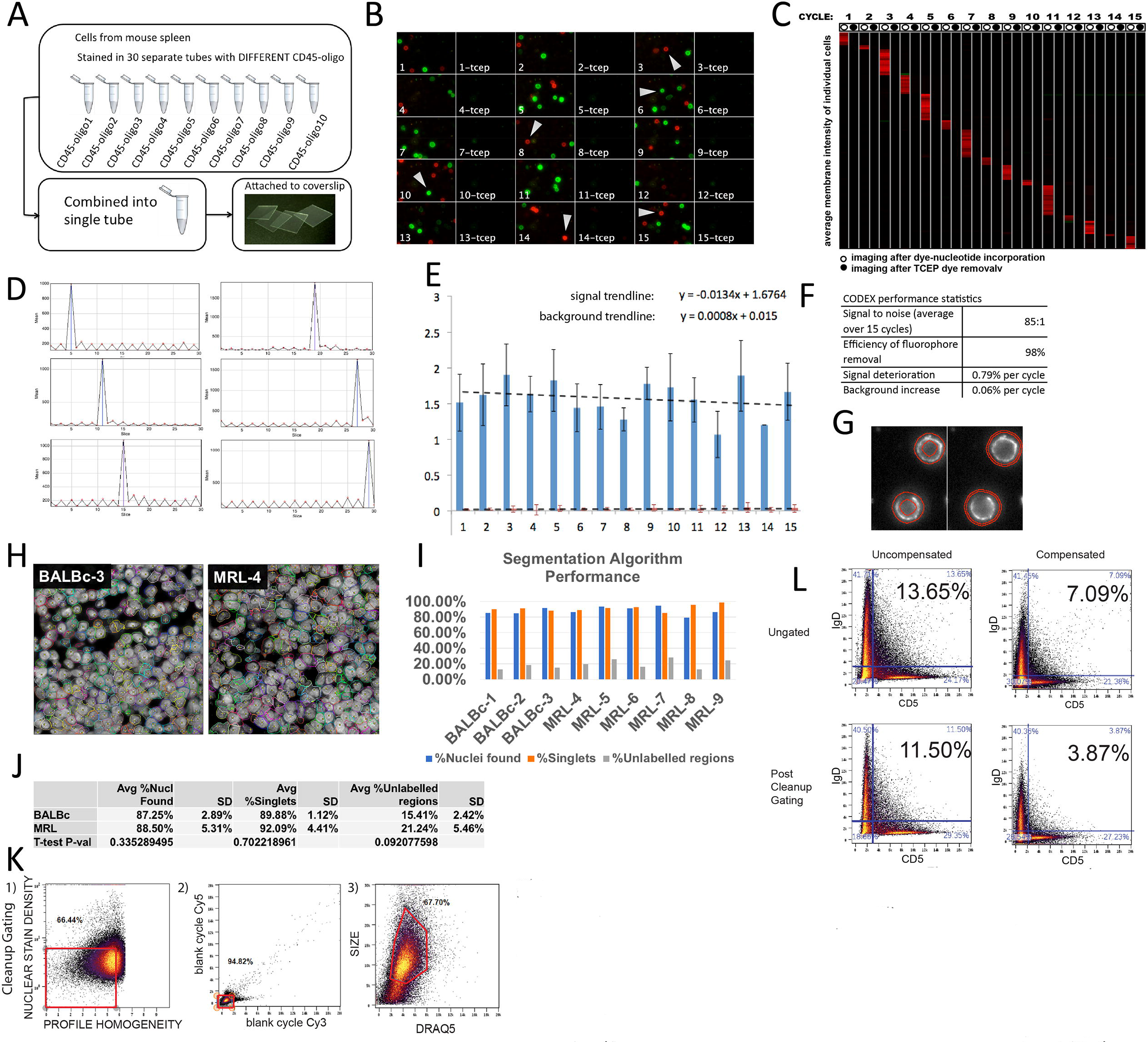
Benchmarking CODEX. Related to Figure 1 and Figure 2. **(A)** Experimental scheme for mimicking the tissue with 30 distinct cell types **(B)** Montage of a fragment of imaging field of the 15 cycles of CODEX used to render the mix of 30 barcoded spleens - first cycle top left last cycle bottom right. **(C)** Heatmap (cycles in columns, cells in rows) showing mean fluorescence per cell membrane for each cell per in each of the 15 CODEX cycles performed on cells of 30 barcoded spleens. Odd columns correspond to imaging after labeled base incorporation. Even columns correspond to imaging after inactivation of staining by TCEP. **(D)** Time-lapse profile of median intensity per cell membrane for individual cells marked by white arrows on (B). **(E)** Average intensity of CD45 antigen expression in “positive” (blue columns) and “negative” (red columns) cells in 15 CODEX cycles of the experiment. (Similar results were obtained for Cy3-positive populations - data not shown). Linear regression was performed to indicate trends in accumulation of background and signal decline associated with cycle number. **(F)** Table summarizing CODEX performance stats. Average signal to noise ratio was estimated from ratio of average signal of all positive cells across all cycles to the signal of all “negative” cells across all cycles. Efficiency of fluorophore removal was estimated from average ratio of ([signal after TCEP in cycle N]- [signal after TCEP in cycle (N-l)])/[signal in cycle N] for cells positive in cycle N across all cycles. Average expected signal deterioration was estimated using the trendline equation from (E). Average background accumulation was estimated by fitting linear trendline into the per cycle ratio of average background to average signal (not shown). **(G)** Image quantification approach used on CODEX data from (A): best focal planes of CODEX stacks were segmented by Cell Profiler. To account for local background the value corresponding to difference between the mean intensity value inside “cell membrane” object (left panel) and the mean intensity inside the external ring object (right panel) was chosen as a representation of the intensity of the antibody signal. In all other experiments custom (see STARS methods) segmantation developed in this study was used **(H)** Sample 500×500 px regions from two samples (BALBc-3 and MRL-4) showing hand-labelled cell centers (yellow crosshairs) and cell outlines detected by the segmentation algorithm (randomly colored). **(I)** Comparison between the hand-labelled cell identification and algorithm-based algorithm identification, expressed in 3 measures: %Nuclei found (how many of the hand-labelled nuclei centers ended up inside the segmented regions), % Singlets (how many of the cell regions with at least one hand-labelled nuclei center contained exactly one cell center) and %Unlabelled regions (how many segmented regions did not contain a hand-labelled cell center) **(J)** Summary statistics comparing the segmentation quality between BALBc and MRL/*lpr* samples. **(K)** Three step cleanup gating strategy based on 1) stain density (nuclear signal divided by cell size) and profile homogeneity (relative variance of signal from cycle to cycle), 2) removing objects with high background by gating on the signal accrued in “blank”(no stain) cycles 3) constraining the cell size. **(L)** Percentage of artefactual double-positive cells in CODEX data from sample BALBc-2 (as seen in the upper right quadrant biaxial flow style plots of mutually exclusive lineage markers IgD and CD5) depending on gating and spill compensation.

**Figure S2.**
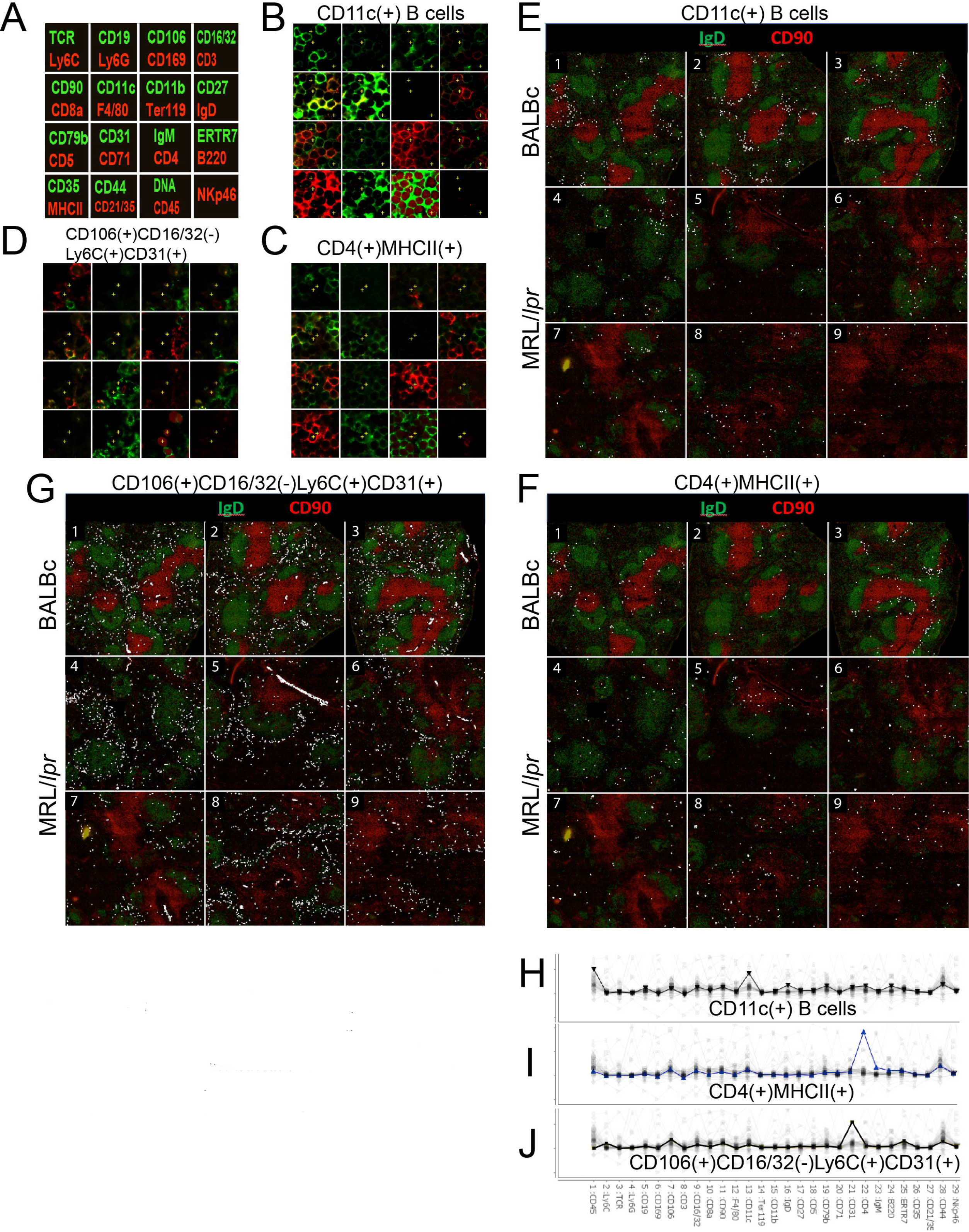
“Cell passports” of selected cell types identified in normal and MRL spleens, Related to Figure 3F and Figure 5B. **(A)** Diagram of per cycle markers for CODEX cycle montages in B,C and D. (**B,E,H**) High resolution montage of CODEX cycles with cells of interest (CDllc(+) B cells) marked with yellow crosses is shown in (B). Low resolution montage of distribution of cells of interest (marked with white circles) in all imaged samples is shown in (E). Average expression profile of all markers in the cells of the selected cell type is shown in (H). (**C,F,I**) Same for CD4(+)MHCII(+) cells. (**D,G,J**) Same for CD106(+)CD16/32(−)Ly6C(+)CD31(+) cells. More examples of “cell passports” can be found in associated online repository (see STAR methods).

**Figure S3.**
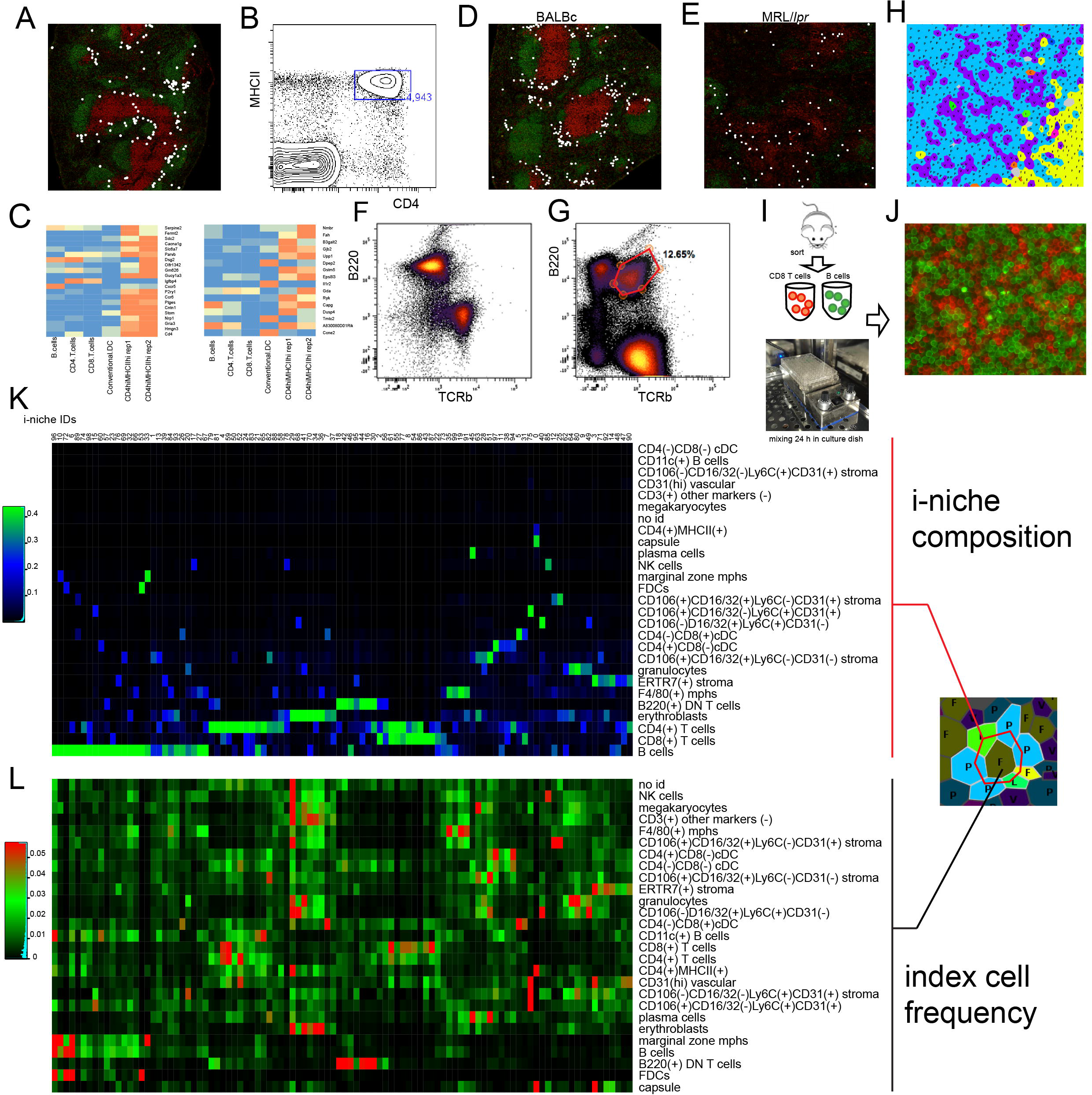
CODEX pinpoints splenic location of unique cell types. Related to Figure 3, Figure 4, Figure 5. **(A)** Distribution of CD4(+)MHCII(+) cells (marked with white circles) in BALBc #2 spleen stained with IgD (green) and CD90 (red) to indicate positions of B and T cells accordingly. **(B)** CD4 and MHCII expression in isolated mouse splenocytes gated negative for all CODEX panel markers and in addition 120g8 (lineage depletion with BD 558451 and dump channel for FITC conjugated or biotinilated antibodies corresponding to the antigens stained with CODEX panel were used for negative gating) except CD4, MHCII, CD45 and CD44. **(C)** CD4(+)MHCII(+) cells within the gate shown in (B) were sorted out and subjected to microarray analysis. CD4 T cells, CD8 T cells, bulk B cells and Conventional CDllc positive dendritic cells were co-sorted as a control. Expression of Lti signature genes (two individual signature sets as inferred in (Robinette et al., 2015)) in sorted cells. **(D and E)** CDllc+ B cells (age associated B cells (ABCs) in normal nd M/lpr spleens. ABCs have been shown to be a key participant in the triggering of certain autoimmune responses (Rubtsova et al., 2017, Rubtsov et al., 2011)) their splenic location has not been previously described in the literature. We observed ABCs to tightly associate with conventional dendritic cells (cDC) and occupy a distinct peri-follicular space in the boundary between PALS and B-zone. Interestingly, these cells diminished in numbers and redistributed towards intra-follicular space in the MRL/*lpr* spleens. **(F and G)** Co-distribution of B220 and TCRb in isolated splenocytes of normal (BALBc) and autoimmune (MRL/*lpr*) mice. Gate in (G) points to significant (^~^13%) presence of B220+ DN T cells in MRL spleen. **(H)** Thread like arrangement of CD8 T cells (purple, annotated with V-letter) has been noticed in PALS of splenic samples across dataset. To examine potential mechanisms driving these structures CD8 Tells and B220 positive B cells were sorted individually from BALBc spleen **(see I)** and later combined in flat bottom microwell plates and mixed at 37C in culture medium. After mixing cells were stained for B220 (green) and CD8a (red) and imaged **(see J)**. Thread like structures similar to what was observed in spleen were detected. **(K)** Heat map showing average frequencies of cell types (rows of heatmap) in the ring of index cell neighbors (see schematics on the right) for all niche clusters (0-99 in columns). **(L)** Heat map shows how different cell types (in rows) are distributed between niches (in columns).

**Figure S4.**
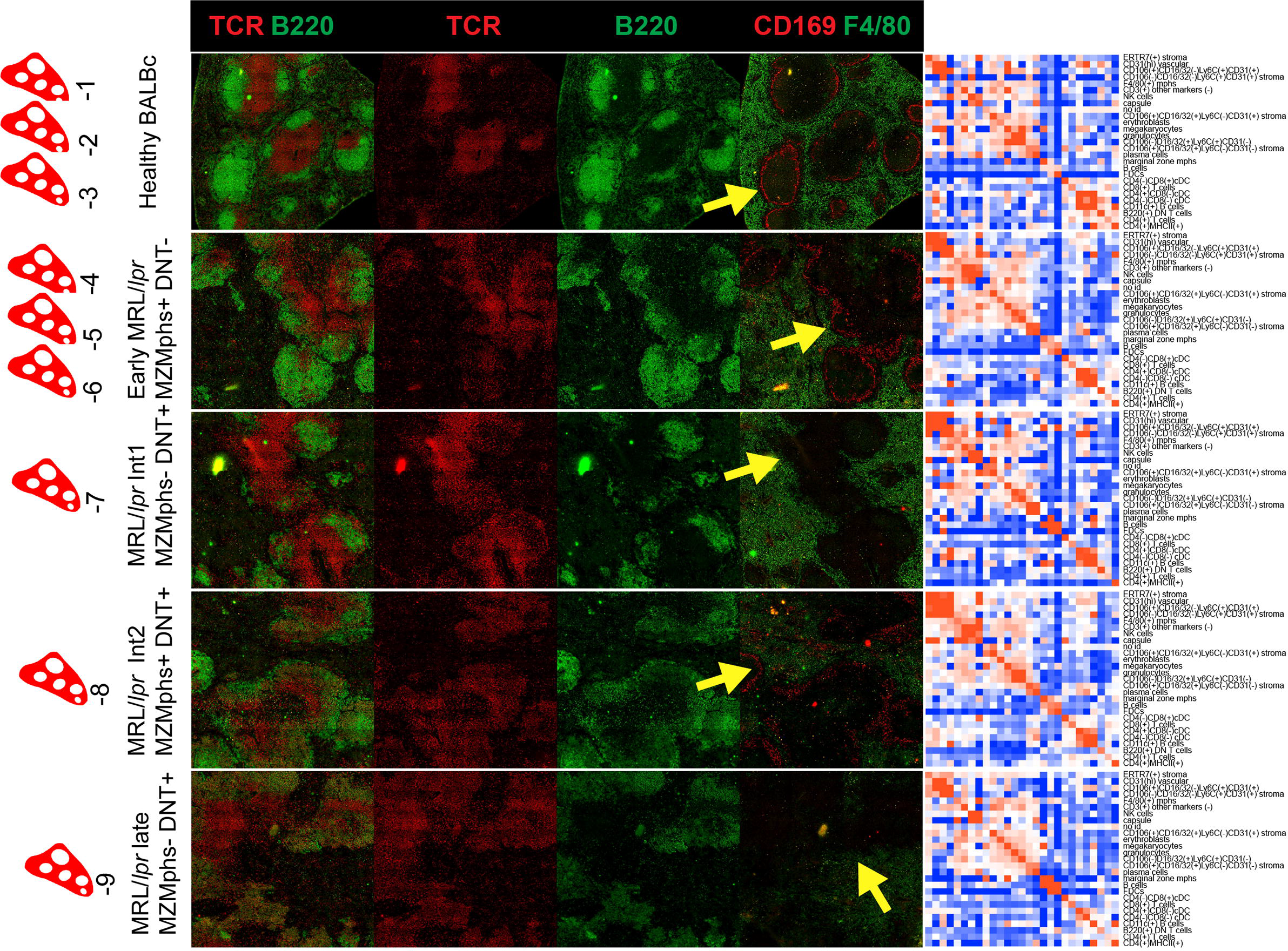
Types of samples in MRL/*lpr* dataset. Related to Figure 5. MRL/*lpr* dataset has 9 samples: 3 control wild type BALBc spleens (BALBc −1,−2,−3 and 6 MRL spleens MRL −4,−5,−6,−7,−8,−9). Based on disintegration of marginal zone as measured by frequency of marginal zone macrophages (MZM’s, - see black asterisk on **Figure 5B** and yellow arrow in this figure pointing to the area where CD169 positive (red) rim of MZMs is expected to be observed) and accumulation of double negative T-cells expressing B220 B cell marker (B220 DN T cells - see red asterisk on **Figure 5B**) MRL spleens were grouped into early (MRL −4,−5,−6), intermediate (MRL −7,−8), and late (MRL −9) types. Early stage was represented by 3 MZM positive DN T cell-low spleens. Two spleens represented the intermediate stage: MZM low DN T cell-low spleen (Int1) and MZM positive DN T cell-positive spleen (Int2). Late stage was represented by single MZM positive DN T cell-positive spleen. A single representative spleen is shown for each stage together with interaction matrix. Color represents odd ratios (observed frequency of interaction/expected frequency of interaction).

**Figure S5.**
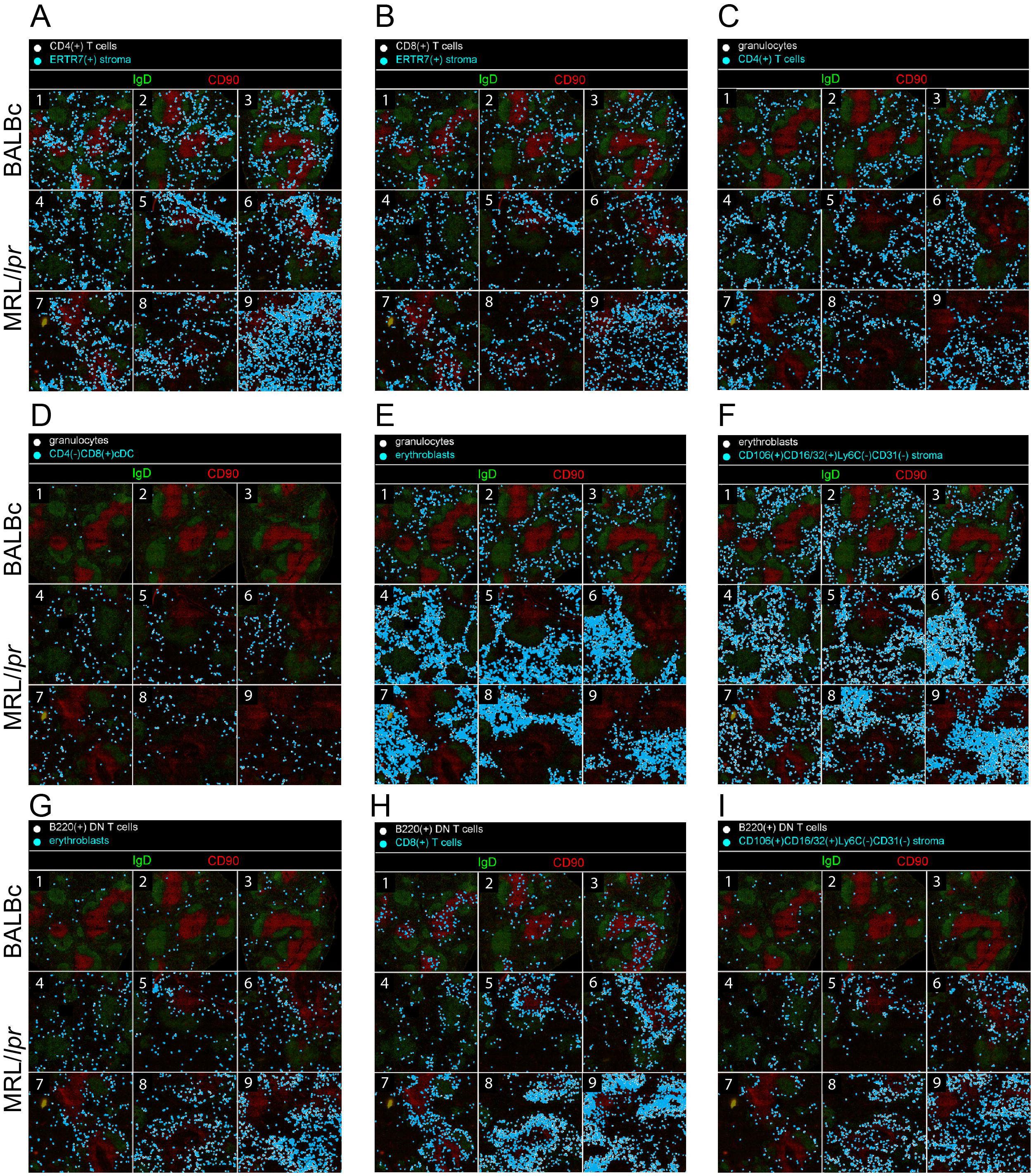
Cross tissue and cross samples distribution of interacting cell pairs for selected types of cell-to-cell interactions, Related to Figure 5C. Interacting cell pairs are marked with white and cyan circles on the montage of IgD CD90 (B cell and T cell markers) staining of every sample of the dataset. Due to cell proximity in most cases cyan circles practically completely overlay white. **(A,B)** Interaction of CD4 and CD8 T-cells with ERTR stroma (change in odds ratio score correlates with change in interaction count). **(C-E)** Interaction of granulocytes with CD4 T cells, dendritic cells and erythroblasts. **(F,G)** Interaction of erythroblasts with stromal and B220(+) DN T cells. Interactions in (C-G) scored as increased in early MRL/*lpr* (−4,−5,−6) as compared to BALBc spleens (FDR of T-test on normalized interaction counts between conditions <0.05, difference in interaction counts>0). **(H,I)** Interactions of B220(+) DN T cells with CD8 T cells and stromal cells. These interactions scored as increased in intermediate and late MRL/lpr (−7,−8,−9) as compared to early MRL/lpr spleens (FDR of T-test on normalized interaction counts between conditions <0.05, difference in interaction counts>0). More examples of cell type pairs with change in interactions across dataset can be found in associated online repository (see STAR methods).

**Figure S6.**
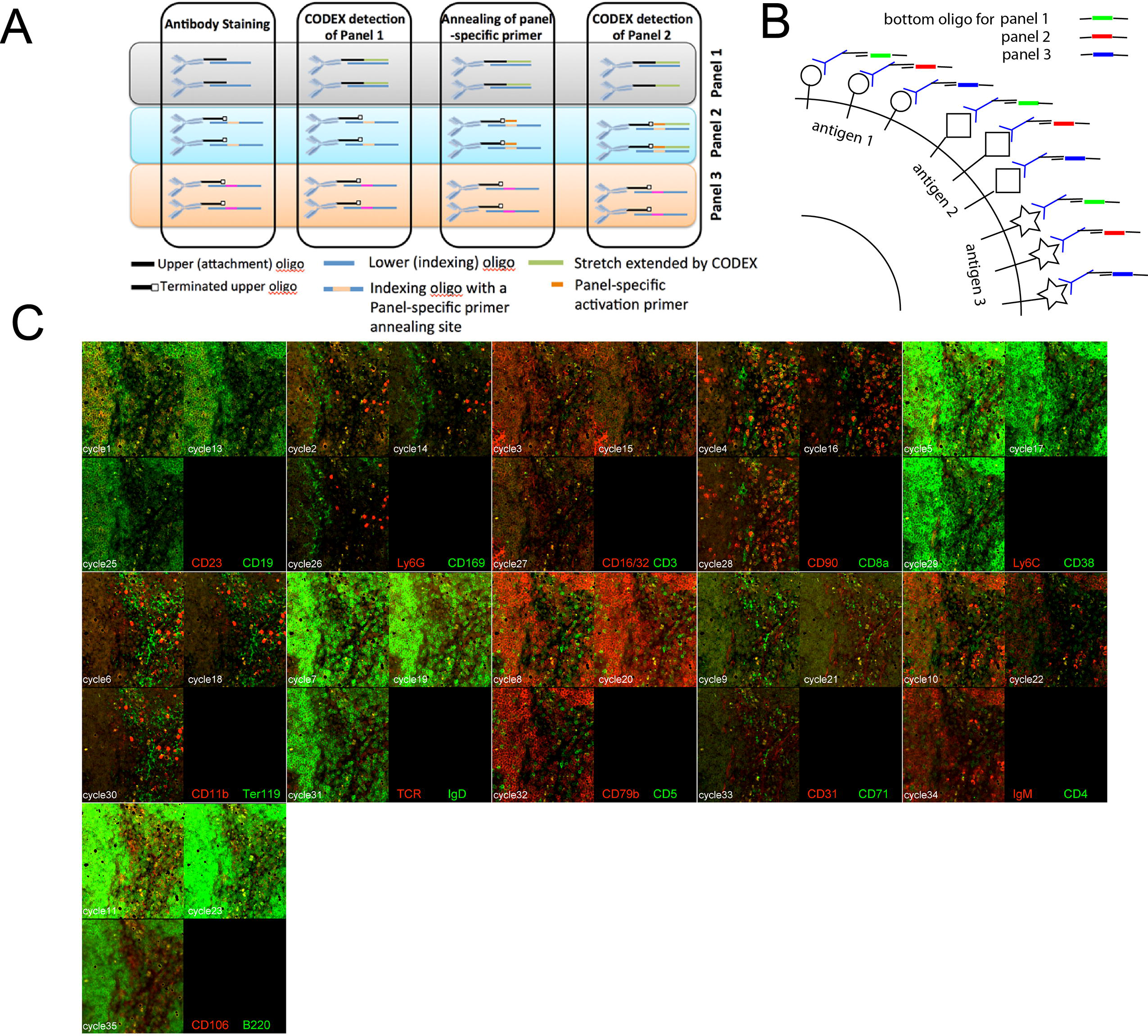
Expanding the multiplexing limit of CODEX by “panels and activators” design, Related to Figure 1A and STAR Methods. **(A)** Diagram of “multipanel”/“activator oligo” CODEX approach. The list of antibodies can be divided in sets such that number of antibodies in each individual set does not exceed the capacity of the multiplexing protocol to render staining without significant signal loss (e.g.30). Each such set of antibodies will be conjugated to “terminated” (the last 3’ base is dideoxy- or propyl- modified) upper strand oligonucleotide of the same sequence as in the original version of the “missing base” approach. The lower strand oligonucleotides will incorporate an additional set-specific region, which will serve as a landing spot for the dedicated primer oligo which is to be on-slide hybridized to the particular subset of the total plurality of the antibodies at the time when they are to be rendered. This approach prevents extension of reads beyond certain threshold and at the same time have an unlimited potential number of antibodies in the sample. **(B)** Schematics of experiment demonstrating the “activator” method and its robustness. Each antigen of a set of 22 surface markers is redundantly detected by three CODEX tag conjugates of the same antibody. The first conjugate is detected during panel 1 rendering, second - during panel 2 etc.‥ Thus the signal for same antigen is detected at different cycles (e.g., 1^st^, 13^th^, and 24^th^) **(C)** Montage of a fragment of imaging field of the 36 cycles of CODEX used to render a mixture of 18 barcoded spleens (similar to design in Figure 2A). Cycles N,N+12 and N+24 all three of which render same pair of antigens are shown per tile for all 11 pairs of antigens (see annotation in the black rectangle of each tile)

**Figure S7.**
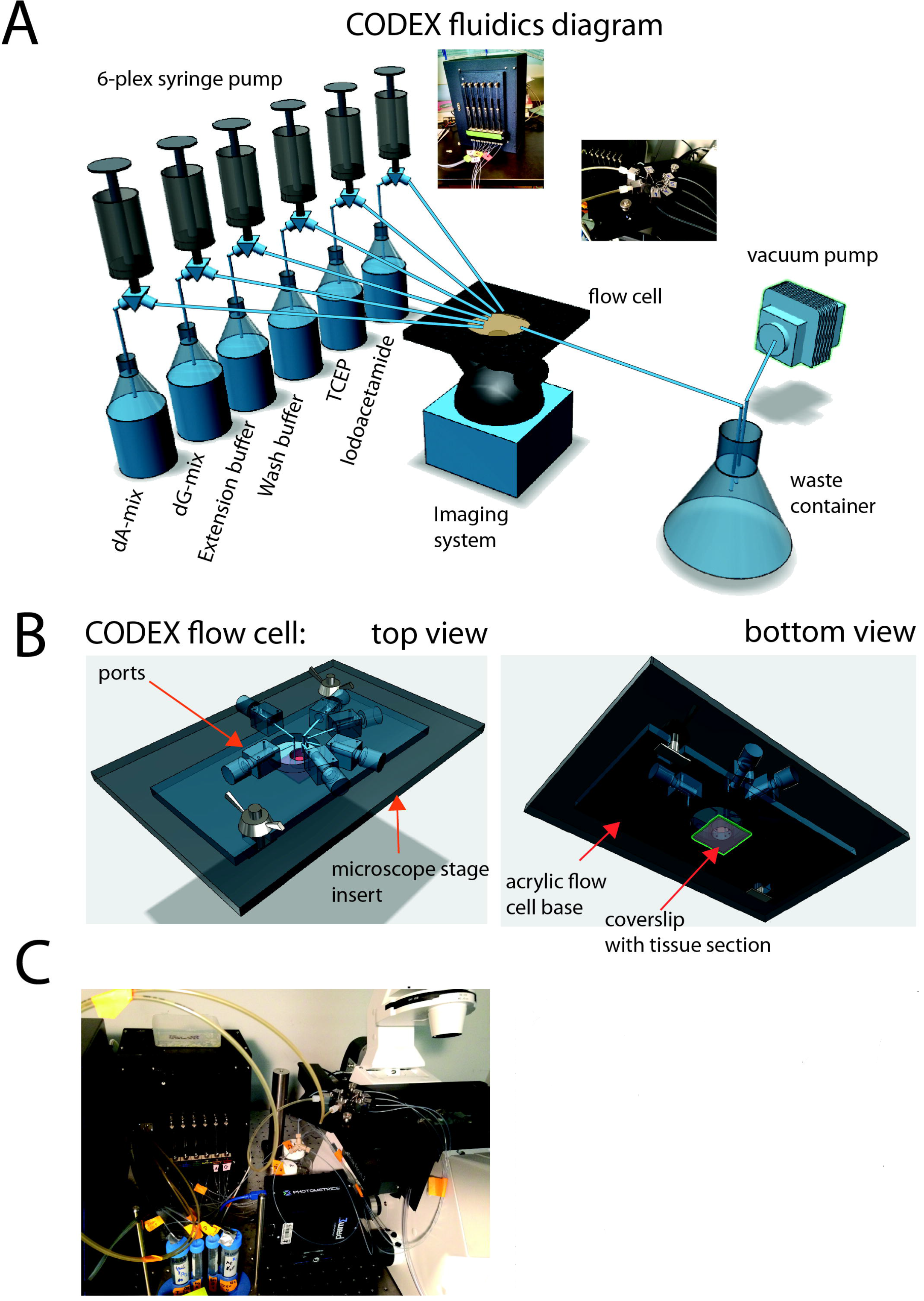
The fluidic setup and stage for running CODEX experiments, Related to STAR Methods. **(A)** General diagram of robotic fluidic setup used in this study. CODEX experiments are done in an open flow cell, which can be imaged in any inverted microscope. Six solutions have to be programmatically delivered and removed from the flow cell, which in the meantime sits in spatially defined position in the imaging system. A combination of 6-channel Tecan syringe pump equipped with 250ul syringes and USB-relay driven vacuum valve was used for iterative solution delivery and removal. Imaging was performed in Keyence BZ-X710 fluorescent microscope configured with 3 fluorescent channels (FITC, Cy3, Cy5) and equipped with Nikon PlanFluor 40x NA 1.3 oil immersion lens. Insets show photographs of actual microscope stage and fluidics robot. **(B)** Detailed 3D model of CODEX stage used in experiments. A metal insert was machined to be compatible with either ASI (Advanced Scientific Instrumentation) or Keyence 3d stages. Disposable (one per experiment) acrylic platform with a circular cutout in the middle was custom designed and lasercut such that it could be attached to the metal stage insert. Before multicycle run the coverslip with a sample was glued to the acrylic base which produced an open flow cell. As opposed to closed, open flow cell design ensures efficient (99.9%) and rapid solution exchange that is critical for CODEX protocol. **(C)** An exemplary photograph of full CODEX setup when attached to an inverted confocal microscope.

## STAR Methods

### Contact for reagent and resource sharing

Further information and requests for resources and reagents should be directed to and will be fulfilled by the Lead Contact, Prof. Dr. Garry Nolan.

### Experimental model and subject details

9 months old female MRL/*lpr* (IMSR Cat# JAX:000485, RRID:IMSR_JAX:000485) (chosen to represent lupus disease at a pronounced splenomegaly stage) and age/sex matched control BALBc mice (IMSR Cat# JAX:000651, RRID:IMSR_JAX:000651) purchased from Jackson Laboratory were used for the study. All animal studies were done in compliance with ethical regulations and procedures set in the Stanford Administrative Panel on Laboratory Animal Care Protocol 15986. In coherence with the primarily technical purpose of the study no animal cohort randomization or investigator blinding to group allocation was performed.

### Experimental method details

#### Oligonucleotide sequences

Single base extension during CODEX can be achieved by either a “missing base” approach (**Figure 1A**) or a “reversible terminator” method (see **Supplementary Movie 1 part 2**). In the case of the “missing base” approach, which was chosen for the experiments outlined in this paper, the top strand of the double-stranded oligonucleotide is covalently bound to the capture agent (in this case, an antibody) and the bottom strand is annealed through hybridization to the top strand. All antibodies contain the same top strand (5’-ATAGCAGTCCAGCCGAACGGTAGCATCTTGCAGAA-3’) and different bottom strands.

The sequence of the bottom strands contains a common region that hybridizes to the top strand (…TTCTGCAAGATGCTACCGTTCGGCTGGAddC-3’) as well as a 5’ variable sequence region that serves as the indexing region. As shown in **Figure 1A**, the overhanging 5’ end of the lower strand of the double-stranded oligonucleotide tag (which forms the overhang) is of the general formula 5’-[C/T]_5_[A/G][5’-C_1-4_/T_1-4_-3’]_n_-TTCTGCAAGATGCTACCGTTCGGCTGGAddC-3’ The first block a short 5-nt stretch of random C/T composition designed to increase the polymerase residence on the DNA duplex. The second block is a single nucleotide (either G or A) that allows for incorporation of a labelled dNTPs (dU-ss-Cy5 or dC-ss-Cy3, respectively). The third block is the “indexing barcode” that consists of *n* random-length homopolymer stretches (1-4 nucleobases each) of alternating “indexing” nucleobases dC and dT that serve as a template for extension of the top oligo with unlabeled nucleotides (dATP and dGTP). Here, *n* specifies the number or extension cycles after which the fluorescent nucleobase will be incorporated into the duplex. Examples of CODEX indexing barcodes are CCCTCC for *n*=3 and CCCTCCTTTCTT for *n*=6. The purpose of having the homopolymer stretches of variable length (e.g. CCCTCCTTTCTT) rather than single base (e.g. CTCTCT) is to increase the polymerase extension specificity and prevent misalignment of upper and lower strands of double-stranded oligonucleotide tags. All oligonucleotide sequences used in this study can be found in **Table S1**.

#### Primer dependent panels to extend the multiplexing capacity of CODEX

CODEX operates using an indexed polymerization step that enables precise incorporation of fluorophores into oligonucleotide-Ab conjugates at predetermined cycles. Although consistent performance of a model antigen (CD45) was observed across 15 cycles of CODEX (**Figure S1A-F**), a gradual accumulation of polymerization errors during each cycle could potentially result in noncognate rendering, and thus diminished and/or non-specific signals at later index cycles. In addition, the use of long single-stranded oligonucleotides that would enable indexing beyond 15 rounds might be problematic due to non-specific binding events to tissues under study.

For the polymerization event to initiate, a 3’ hydroxyl is required. Thus, we reasoned that dedicated primers (each containing a distinct initiating sequence with a 3’ hydroxyl) could be used to activate distinct subpanels of antibodies (**Figure S6A**). This would allow design of antibody panels exceeding 30 markers into subpanels, each with a subpanel-specific activation sequence designed 5’ to the indexing region. In this design, the antibody attachment linker is terminated with ddC, such that the extension is only possible after a hybridization of a hydroxyl-containing panel-specific activation primer.

The feasibility of such multipanel CODEX design and the robustness of CODEX protocol after many cycles and its independence of staining from the cycle number were tested in a model experiment. A 22-color panel of antibodies (11 cycles) conjugated to a terminated top oligonucleotide, was hybridized with lower oligonucleotides of 1^st^, 2^nd^, and 3^rd^ panels (**Figure S6B**). Thus, every antigen is detected thrice by the same antibody conjugated to oligonucleotides of 3 different panels. Each panel can only be rendered after annealing of a panel-specific activator oligonucleotide. The staining was rendered in 36 cycles (11 detection cycles + 1 blank no-antibody cycle per activator oligo) of CODEX with additional activator oligonucleotide hybridization step between each of the 3 panels. The signal for same antibody detected at different cycles (e.g., 1^st^, 13^th^, and 24^th^) was consistent across the three panels (**Figure S6C**). This panel-activator design extends CODEX to a theoretically unlimited multiplexing capacity, bounded only by the speed and resolution of the imaging process itself and the time required for each imaging cycle.

#### CyTOF CODEX comparison

Cell preparation and staining by metal tagged antibody for CyTOF analysis was performed as described before (Spitzer et al., 2017). Mass cytometry was performed on a CyTOF™ 2 mass cytometer (Fluidigm) equilibrated with ddH20. For CODEX analysis, isolated spleen cells were stained a panel of antibodies conjugated to indexing oligonucleotides. Samples were fixed to a coverslip (**Figure 2A**) and imaged over 12 cycles of CODEX protocol. Images were segmented using the *in situ* cytometry software toolkit developed for this study (see **Figure 2A** for exemplary segmentation of the cell spread), and the staining of individual cells across the indexing cycles was quantified. Segmentation data was converted into flow cytometry standard (FCS) format and analyzed using the conventional flow cytometry analysis software Cytobank.

#### Antibody conjugation, staining and CODEX rendering

Detailed step-wise CODEX protocols can be found online (see Key Resource Table and Data and Software availability section below). For full list of antibody clones and vendors see **Table S1**. Custom manufactured microfluidic setup (**Figure S7 A-C**) was used to automate CODEX solution exchange and image acquisition. Instrument and blueprints and control software are available upon request.

### Primer dependent panels

Rendering of antibodies with spacers followed the same procedure as the standard CODEX protocol with the exception of the following differences. Before proceeding to rendering next spacer dependent panel, the stained cells were incubated with a spacer oligonucleotide (1μs;M final concentration in buffer 405) at room temperature for 10 minutes. Cells were washed 4X with buffer 4 and rendering proceeded as usual. To initiate each additional spacer set, the spacer incubation step was repeated using corresponding spacer samples.

### Imaging

Images were collected using a Keyence BZ-X710 fluorescent microscope configured with 3 fluorescent channels (FITC, Cy3, Cy5) and equipped with Nikon PlanFluor 40x NA 1.3 oil immersion lens. Imaging and washes were iteratively performed automatically using a specially developed fluidics setup (**Figure S7 A-C**). Images were subject to deconvolution using Microvolution software (www.microvolution.com). The staining patterns of 28 DNA-conjugated antibodies were acquired over 14 cycles of CODEX imaging and overlaid with 2 additional fluorescent antibodies, CD45-FITC and NKp46-Pacific Blue and a DRAQ5 nuclear stain (**Figure 3A** and low-resolution views in **Supplementary Movie 2**). Each tissue was imaged with a 40x oil immersion objective in a 7×9 tiled acquisition at 1386×1008 pixels per tile and 188 nm/pixel resolution and 11 z-planes per tile (axial resolution 900 nm). Images were subjected to deconvolution to remove out-of-focus light. After drift-compensation and stitching, we obtained a total of 9 images (one per tissue) with x=9702 y=9072 z=ll dimensions, each consisting of 31 channels (30 antibodies and 1 nuclear stain).

### QUANTIFICATION AND STATISTICAL ANALYSIS

#### Initial image analysis and segmentation

For each imaging field analyzed by CODEX multidimensional staining multi-color z-stacks collected during individual cycles were aligned against reference channel (CD45) by 3D drift compensation (Parslow et al., 2014). If necessary individual fields covering large tiled areas were “stitched” using dedicated ImageJ plugin (Preibisch et al., 2009). For the 22-channel experiment on dissociated cells attached to coverslip (**Figure 2**) images corresponding to the best focal plane of vertical image stacks collected at each acquisition step of CODEX were chosen for quantification. For the 31-channel main experiments on mouse spleen sections, the segmentation was performed on the whole image stack using a volumetric segmentation algorithm described below. For this study, we purposefully developed a 3D image segmentation that combines information from the nuclear staining and a ubiquitous membrane marker (in this case CD45) to define single-cell boundaries in crowded images such as lymphoid tissues. This algorithm inverts the membrane image and multiplies it with the nuclear image, creating a synthetic image with enhanced contrast between neighboring nuclei. This image is subject to low-pass FFT filtering and an then individual cell objects (collections of voxels) are identified using a gradient-tracing watershed algorithm. Per-cell intensities were quantified by integrating the intensity of each channel within a given cell object and divided by the region size in voxels.

We benchmarked the segmentation algorithm against a dataset of BALBc mouse spleen images with expert hand-labelled nuclei in and we found the algorithm was able to correctly identify 87.25%±2.89% cells, of which 89.88%±1.12% were singlets (one-to-one correspondence between a hand-labelled cell and a segmented object) (**Figure S1 H-J**). For each segmented object (i.e., cell) a marker expression profile, as well as the identities of the nearby neighbors were recorded using Delaunay triangulation (http://dx.doi.Org/10.17632/zjnpwh8m5b.l).

#### Spatial spillover compensation

Accurate segmentation per se is not sufficient to obtain high-quality estimate of single-cell expression from images. The reason for that is that in lymphoid tissue the cells are so tightly adjacent that their membrane signals can partially overlap, resulting in blending of signals between neighboring cells, the phenomenon termed spatial spillover. In order to compensate for that, we estimated the cell-to-cell signal spill coefficients based on the fraction of shared boundary between each pair of cell objects, resulting in a banded matrix (most cells don’t have any shared boundaries). To compensate the cell-to-cell spill, the raw intensity vector was multiplied by the inverse spill matrix (Figure 2D).

Besides the spatial signal spillover, there are other factors that add to artefactual cell-like objects: debris misidentified as cells, doublets (two adjacent cells merged together) as well as autofluorescent objects, both of which can lead to spurious as double-positive signals on the biaxial scatterplots. By analogy to how debris and doublets are eliminated from FACS data by applying special ‘Singlet’ gates to SSC-FSC parameters, we devised a ‘cleanup’ gating strategy based on several quality control parameters: nuclear stain density (nuclear signal divided by cell size), profile homogeneity (relative variance of signal from cycle to cycle), background staining on blank cycles and, finally, nuclear signal and cell size (Figure S1K). We found that applying those filtering gates had a synergetic effect with the compensation, reducing the frequency of spurious double-positive cell signals by approximately an additional factor of 2 (**Figure S1L** e.g. compare fraction of CD5/IgD double positive cells in Ungated-Compensated and Post Cleanup gated - Compensated in (L)).

#### Cell type definition

The 9-spleen dataset was subject segmentation, quantification, compensation and cleanup gating, as described above, yielding a total of 734101 30-dimensional single-cell protein marker expression profiles (**Figure 3C**, **http://dx.doi.Org/10.17632/zjnpwh8m5b.l**). The segmented CODEX data was subject to automated phenotype mapping algorithm X-shift that was previously developed and validated on CyTOF data (Samusik et al., 2016) (**Figure 3C**). 58 phenotypic clusters inferred by X-shift clustering were manually annotated (**Figure 3C, D** and **Table S2**) based on the 30-color marker expression profile and thorough visual inspection of the representative image samples (**Figure S2A-J**, more examples of “cell passports” can be found in associated online repository - see STAR methods). Some clusters were found to originate from imaging artifacts such as dust and tissue sectioning defects. That reduced the overall number of cell-like objects to 707466. Each cluster was assigned to one of 27 broadly defined single-cell phenotypic groups (cell types), which in some cases could be clearly matched to major immune cell types and in others were named according to expression of distinguishing surface markers (see cluster annotation and cell counts in **Table S2**).

#### Cell interaction analysis

To define for each cell the neighbors of the first (immediate) tier of proximity Delaunay graph was computed for the dataset (**http://dx.doi.Org/10.17632/zjnpwh8m5b.l**). The odds ratio of cooccurrence of cell type A and cell type B was estimated as the observed frequency of co-occurrence (mean of the beta-distribution, with parameter alpha = number of edges connecting cell types A and B and parameter beta = total number of edges minus number of edges connecting A-B) divided by the theoretical frequency of co-occurrence (total frequency of edges incident to type A multiplied by the total frequency of edges incident to type B) see **Table S3**. The odds ratios are represented in heatmaps on **Figure 3G**, with a range of values from less than 1 to more than 1 meaning that two cell types are, respectively, less or more likely to co-occur than expected by chance. The significance of the difference from zero was tested using binomial distribution (probability of getting an observed number of interactions between A and B (successes) amongst the total number of registered interactions (number of trials) given the theoretical probability of A-B interaction (probability of success)).

The significance of change of interaction frequencies or log-odds ratios were computed between BALB/c and Stage 1 (early) MRL using pairwise T-test. However, the same procedure could not be applied to testing BALB/c versus MRL/*lpr* Stages 2 and 3 because of high sample-specific variation in those more advanced disease stages. Therefore we scored computed the deviation of those Stage2/3 values from BALB/c using χ^2^ statistics because it does not require Stage 2/3 samples to have a common mean.

The P-values were subject to FDR correction using Benjamini-Hochberg procedure. Interactions that were considered significant for FDR q-value < 0.05 or > 0.95 (**Table S3**).

In order to estimate the overall deviation of either interaction frequency matrices or log-odds ratio matrices, the matrices were subject to z-transformation based on the mean and the SDs of the BALB/c samples, and then χ statistics was computed as square root of the sum of squares of all elements of the z-score transformed matrices (**Figure 5F**).

#### i-niche analysis

The i-niche analysis was performed based on 2-dimensional Delaunay triangulation of the cell center coordinates. Delaunay triangulation and the related concepts of Voronoi Tesselation and Gabriel graphs were previously applied in eco-geographical analysis of species distribution (Gabriel and Sokal, 1969) and therefore were deemed as equally applicable to the analysis of tissue organization on the single-cell level. The i-niche is defined as a set of first-order Delaunay neighbors of the given ‘index cell’, i.e. the i-niche cells are the ones that are directly connected to the i-cell with edges in the Delaunay triangulation of cell centers. We distinguish i-niche from the more formal understanding of “niche”, which is often used in stem cell literature and where numbers of cells in the niche and their placement within the niche is undefined. In our definition, we allow the central cell to be of any type and are counting the cell types present in the ring. This flexible definition allows for multi-cellular interactions around a central cell to define the biology of that cell (and vice versa). Computationally, the i-niche window slides from cell to cell, considering each set of adjoining cells—and therefore allows consideration of the constituencies of different central cell types that might populate a given i-niche. We understand that our current definition is arbitrary and could be extended to include other specific cell arrangements—including, though beyond the scope of the current work, a 3D sphere of cells contacting the index cell.

#### Neural network training and data analysis

##### Preprocessing

Image stacks were maximum-intensity projected following deconvolution. Data was quantile normalized to 4 levels (0, 0.25, 0.5 and 0.75 quantiles). A baseline model was able to distinguish models without this discretization and normalization, suggesting strain-specific differences in antibody staining intensity.

##### Training and cross validation split

Four spleen samples (two BALB/c and two MRL/*lpr*) were chosen as training samples. The remaining five spleens tissue samples (one BALB/c and four MRL/*lpr*) were used for testing the trained model. For cross-validation, different combinations of spleens were allocated to training and test sets. During training, 224×224 images were randomly extracted from the training tissue samples, at 1x, 0.5x and 2x zoom. At 1x zoom, there would be 6804 non-overlapping image patches in the training dataset. The trained models were tested on 4500 patches, at 1x zoom. Hyperparameters were manually tuned on 500 randomly selected images from the testing spleens. The Adam optimizer was used for training with an initial learning rate of 0.0001.

###### Baseline model

A logistic regression model was trained by averaging marker intensities across the image. L2 regularization was used for weights.

###### Neural network architecture

To avoid the learning of trivial sample-specific staining variation, data were quantile normalized sample-wise and each marker was discretized to four levels. Since disease-specific hallmarks could potentially be present at multiple scales, the training data for our neural network was extracted at multiple levels of magnification. A simple regularized logistic regression model that considered only average marker expression and did not incorporate spatial information was unable to successfully distinguish patches normal and MRL/*lpr* spleens, whereas the trained neural network model consistently achieved a 90% precision of classification of image patches during cross-validation.

A fully convolutional network architecture was used, with the following layers. To generate a prediction for an entire image patch, a global max-pooling layer was used.

1. Conv3 60
2. Conv3 120
3. Conv3 64
4. Batch Norm
5. Conv3, 64
6. Max pooling 2×2
7. Conv3,128
8. Conv3,128
9. Max pooling 2×2
10. Conv3, 256,
11. Conv3,256
12. Conv3,256
13. Max pooling 2×2
14. Conv3,512
15. Conv3,512
16. Conv3,512
17. Convl,256
18. Convl,64
19. Convl,l
20. Global max pooling
21. Sigmoid

Weights for layers 5-16 were initialized from the VGG-16 pretrained model. The model was trained with cross-entropy loss.

##### Regularization

L2 regularization (0.1) was used for network weights. L1 regularization was applied to the feature map output after layer 19 to encourage sparse activations

##### Whole sample activations for test set

Since the network was fully convolutional, it could be applied to images of any dimension. The network was applied to entire fields of view individually. The activation maps were obtained as the output after layer 21.

##### Aligning cell type information

Each cell was assigned the MRL/*lpr* score of the corresponding pixel in the image.

##### Enrichment and neighborhood analysis

FDR controlled chi-squared tests of proportions were carried out to determine enrichment of specific cell types in the top 10% of cells by MRL/*lpr* score. For neighborhood analysis of dendritic cells, the composition of the neighborhoods (cell centers within 30 pixels) of the top 300 cells (by MRL/*lpr* score) were compared to the composition of the neighborhoods of the bottom 300 cells. Only cells with positive neural network assigned MRL/*lpr* score, in MRL/*lpr* regions, were considered for this analysis.

#### Data and software availability

Software used in the paper for parsing image data can be obtained at: https://github.com/nolanlab/CODEX

Data tables can be downloaded from Mendeley: https://dx.doi.Org/10.17632/zjnpwh8m5b.l

All primary image data, high resolution focused montages, complete single cell data tables and various additional information can be obtained at: https://welikesharingdata.blob.core.windows.net/forshare/index.html

Flow formatted segmented data can obtained from online repository page (link above) or from Cytobank: CODEX on spreads of isolated mouse splenocytes (Figure 2B):
https://communitv.cvtobank.org/cvtobank/experiments/69534 https://flowrepository.org/experiments/1686

CyTOF on isolated splenocytes (Figure 2B): https://community.cvtobank.org/cvtobank/experiments/69533 https://flowrepository.org/experiments/1687

CODEX on BALBc spleen tissue sections (Figure 2E): https://community.cvtobank.org/cvtobank/experiments/69889 https://flowrepository.org/experiments/1688

## Supplementary Table Legends

**Table S1. List of CODEX antibodies and oligonucleotides, Related to Figure 2 and Figure 3**.

Excel file with four spreadsheets corresponding to multidimensional staining experiments performed in the study **(CODEX panel for cell spreads)** List of 24 antibodies (23 DNA conjugated + CD45 FITC for counterstain), upper and lower nucleotides used for CODEX staining of isolated splenocytes. **(CODEX panel for spleen tissue)** List of 30 antibodies (28 DNA conjugated + CD45 FITC and NKp46 PacBlue), upper and lower nucleotides used for comparative CODEX staining of normal BALBc and lupus afflicted MRL/*lpr* spleen sections. **(CYTOF panel for spleen cells)** List of 23 metal conjugated antibodies antibodies used in CyTOF analysis of isolated splenocytes. **(Activator driven CODEX panels)** List of 22 antibodies (22 DNA conjugated + CD45 FITC for counterstain), upper, lower and activator nucleotides used for activator driven CODEX staining of isolated splenocytes (see exp. Schematics in Figure S12).

**Table S2. X-shift cluster annotations and cell counts, Related to Figure 3**

Excel file with 58 clusters identified by X-shift analysis, their annotations and resulting across dataset counts for 27 imaging phenotypes identified in this study

**Table S3. Dynamics of average cell type to cell type interaction frequency and strength across dataset, Related to Figure 3G.**

Excel table with three spread sheets. **Full data** contains odds ratios; direct counts of interactions as well as various differential metrics for comparisons off frequency and strength of cell type to cell type interactions between early MRL and control (BALBc) and intermediate-late MRL and early MRL. **Early vs control** shows top candidate cell type pairs selected based on the change in strength (odds ratios) or frequency of interactions between early MRL spleen and control spleens. **Late vs early** shows top candidate cell type pairs selected based on the change in strength (odds ratios) or frequency of interactions between combined intermediate and late MRL spleens and early MRL spleens.

**Table S4. Linear regression model for marker expression level based on niche and cell type shows importance of niche, Related to Figure 4D,E**.

The overall role of the niche in defining marker expression was evaluated by constructing a linear regression model of marker expression with cell type identity and niche as two feature variables. This Excel file shows F and P values for the contribution of niche to the model. The F value is the ratio of the mean regression sum of squares for the model including just cell type to the full model including both niche and the cell type. Its value ranges zero to an arbitrarily large number. A larger F value suggests that the niche has a larger contribution in explaining the variance observed in the expression levels of each marker. The value of Pr(>F) is the p-value against the null hypothesis that including the niche in the model does not improve the fit.

